# Multiomic Analysis Reveals an IFN-driven Cellular Landscape Effectively Targeted by Ruxolitinib in Hailey-Hailey Disease

**DOI:** 10.64898/2026.06.01.729262

**Authors:** Sara Ceccacci, Hoang Duc Minh Pham, Marion Dessaignes, Jasmin Mirzayan, Kevin Roger, Lam C. Tsoi, Paul W. Harms, Johann E. Gudjonsson, Maria Emilia Puig Lombardi, Ida Chiara Guerrera, Alain Hovnanian

## Abstract

Hailey-Hailey disease (HHD) is a rare autosomal dominant genodermatosis characterized by skin blistering and erosions in intertriginous regions, frequently complicated by secondary infections leading to substantial impairment in quality of life. No targeted mechanism-based therapies are currently available. Here, we applied a multiomics approach to define the molecular and cellular landscape of HHD. Bulk transcriptomics and proteomics uncovered a striking interferon (IFN) signature in HHD skin lesions. Single cell and spatial transcriptomics analyses revealed inflammatory niches, where immune, epithelial, vascular and stromal cells create a multi-compartment IFN-driven signaling network, that sustains a feed-forward amplification loop essential for chronic inflammation. Crucially, in nine patients with refractory HHD, topical treatment with the JAK1/2 inhibitor ruxolitinib led to rapid and durable re-epithelialization with drastic reduction in pain, itching, oozing and skin inflammation, significantly improving patient quality of life. Collectively, our findings identify IFN signaling as a key pathogenic driver in HHD and support topical JAK inhibition as an effective therapy, redefining the standard of care for individuals living with HHD.

## INTRODUCTION

Hailey-Hailey disease (HHD), also known as familial benign chronic pemphigus, is a rare autosomal dominant genodermatosis, characterized by impaired epidermal keratinocyte adhesion (acantholysis)^1^. The incidence of HHD is estimated at 1 per 50’000 individuals, with no gender predilection^2^. Clinically HHD manifests as recurrent vesicopustules and blisters, weeping erosions with fissures and rhagades in intertriginous or friction-prone regions. The lesions often expand in annular plaques with peripheral crusty borders^3^. They are frequently complicated by infections that exacerbate inflammation, itching and pain, substantially compromising patients’ quality of life (QoL). The course is chronic with flare-ups that could be triggered by heat, sun exposure, friction, sweating, and stress. Histologically, acantholysis gives the epidermis a characteristic “dilapidated brick wall” appearance, often accompanied by epidermal hyperplasia, parakeratosis and dermal perivascular lymphocytic infiltrate^1^.

HHD is caused by pathogenic variants in *ATP2C1*, which encodes the Secretory Pathway Ca²⁺-Transporting ATPase type 1 (SPCA1) that actively transports Ca²⁺ and Mn²⁺ from the cytosol to the lumen of the Golgi apparatus ^4,5^. SPCA1 dysfunction leads to reduced Golgi Ca²⁺ levels, which may impair intraluminal Golgi processes, including post-translational modifications, trafficking and sorting of molecules crucial to cell-cell adhesion, such as desmosomal proteins ^6,7^.

Current treatments for HHD are non-specific and primarily aim to dampen inflammation and prevent secondary infections. They mainly rely on potent topical corticosteroids and/or antimicrobials for mild forms, escalating to oral antibiotics, ultra-potent topical corticosteroids or topical calcineurin inhibitors in case of recurrent exacerbations^8^; HHD is often refractory to treatments. For severe forms, invasive procedures (botulinum toxin type A injections to reduce sweating, laser therapies, dermabrasion, surgical excision of the affected area) or systemic immunosuppressive agents (methotrexate, cyclosporin, apremilast) ^9–11^ have been used. Low doses of naltrexone, a long-acting opioid receptor antagonist, have shown variable efficacy^12^. Recent case reports have described promising results in HHD refractory forms treated with subcutaneous injections of dupilumab^13–15^, an IgG4 monoclonal antibody targeting IL-4Rα, as well as with oral administration of JAK inhibitors^16–18^. Notably, two newly published case reports documented significant improvement with the topical JAK1/2 inhibitor ruxolitinib^19,20^. However, the mechanism of action of JAK inhibitors in HHD has not been investigated to date.

Here, we applied a multiomics strategy to define the key molecular signatures of HHD lesions and to pinpoint the main cellular drivers within their native tissue context^21^. Specifically, we performed bulk transcriptomic and proteomic analyses on skin biopsies from a cohort of patients with HHD, while single-cell and spatial transcriptomic analyses (Visium and Xenium) were conducted on a smaller subset. Collectively, these complementary omics layers provide an unprecedented, high-resolution molecular and cellular landscape of the HHD pathogenic microenvironment, directly leading to a targeted therapy which has proven successful on nine patients treated.

## RESULTS

### 1. Integrated bulk transcriptomics, proteomics and spatial analyses uncover IFN-driven immune response in Hailey-Hailey disease

To define a distinctive global proteogenomic signature in HHD patients, we integrated bulk transcriptomic analysis of full-thickness skin biopsies (23 healthy controls and 20 patients) with total proteomic analysis (24 patients, 15 in common with the transcriptomic cohort) (**Supplementary Table S1, see *Patient and healthy control biopsies* in Methods**). For each patient, skin biopsies were collected from lesional (L) and non-lesional (NL) areas.

Across the bulk transcriptomics and proteomics datasets, we quantified 19,580 transcripts and 8,191 proteins, of which 3,610 transcripts^(t)^ and 1,216 proteins^(p)^ were significantly dysregulated in L *versus* NL samples (FDR < 0.05, |log2FC| > 1) (**Supplementary Table S2**). Over-representation analysis of significantly upregulated transcripts and proteins showed common enriched pathways mainly pointing to: i) innate immune response, ii) T-cell receptor (TCR) activation and regulation, iii) IFN and IL-4/IL-13 signaling (**Figure 1A, Supplementary Table S3**).

**Figure 1.**
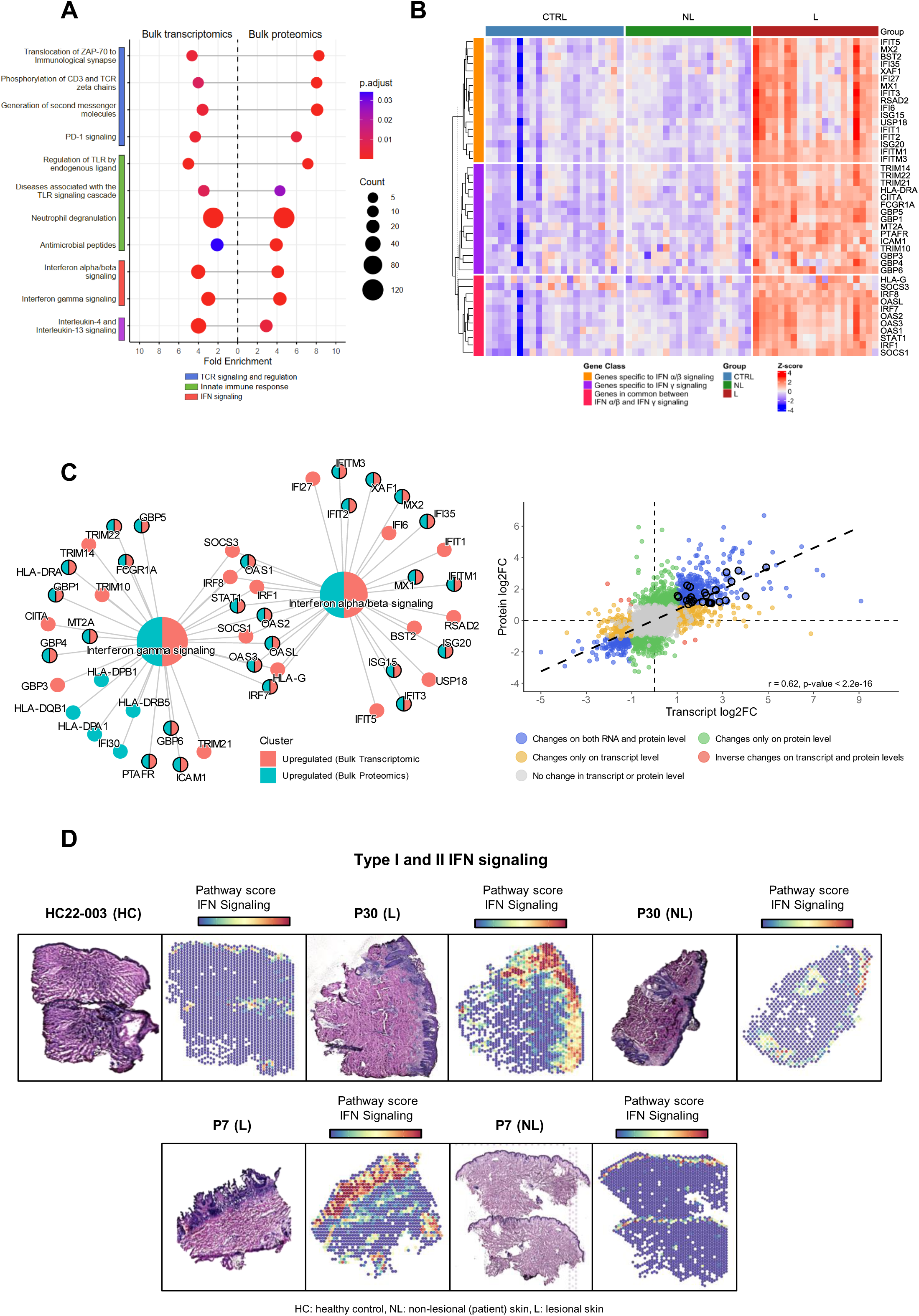
Integrated transcriptomic, proteomic and spatial analyses reveal upregulation of IFN signaling in HHD lesional skin. **(A)** Over-representation analysis of significantly upregulated transcripts and proteins in lesional (L) vs non-lesional (NL) samples, highlighting common enriched pathways. Circle size indicates the number of genes/proteins contributing to enrichment; color reflects adjusted p-value. **(B)** Heatmap of IFN-stimulated genes (ISGs) across control (CTRL), non-lesional (NL), and lesional (L) skin samples. Z-scores represent normalized expression values. Genes are annotated as IFNγ-specific, IFNα β-specific, or shared ISGs. **(C)** Left: Network representation of significantly upregulated genes and proteins (L vs NL) associated with interferon signaling. Node color distinguishes transcriptomic vs proteomic upregulation. Right: Correlation of transcriptome and proteome changes (log2FC) across samples. Colored points represent concordant or discordant regulation. IFN genes are circled in black. **(D)** Spatial transcriptomics maps (VisiumSD assays) showing IFN signaling pathway activity scores in representative healthy control (HC), patient non-lesional and lesional skin sections.

The activation of the innate immune response is reflected in our data by the upregulation of the Toll-like receptor (TLR) signaling (TLR1^t^, TLR2^tp^, TLR7^t^, TLR10^t^, CD14^t^, LY96^t^, MYD88^t^), neutrophil degranulation, and antimicrobial peptides (DEFA1^p^, DEFA3^p^, CAMP^tp^, BPI^p^). We also observed activation of the adaptive immune response, primarily mediated by T cells, with upregulation of key components of the TCR signaling pathway (CD3D^t^, CD3E^t^, CD3G^tp^, CD247^t^, LCK^tp^, ZAP-70^tp^, GRAP2^p^, LCP2^tp^, ITK^t^).

Consistent with increased TLR and TCR signaling^22^, our data revealed significant upregulation of type I (α/β) and type II (γ) IFN pathways. As shown in the heatmap in **Figure 1B**, lesional samples displayed a strong IFN signature compared with healthy control and non-lesional samples. Remarkably, around 30 IFN-related genes were found upregulated in lesional *versus* non-lesional samples at the transcriptomic and proteomic levels (**Figure 1C Left**). Gene quantification across the two omics showed a moderately high Pearson correlation (r=0.62, p-value<2.2e^-16^), suggesting a parallel regulation of the IFN pathway (**Figure 1C Right**).

Specifically, we observed upregulated expression of key components of both type I and type II IFN pathways, supporting activation and modulation of the canonical IFN downstream JAK/STAT signaling axis^23^, as evidenced by increased STAT1^tp^ together with induction of its negative regulators^24^ SOCS1^t^ and SOCS3^t^. Interferon regulatory transcription factors (IRF1^t,^ IRF7^tp^, IRF8^t^) and IFN-stimulated OAS proteins^25^ (OAS1^tp^, OAS2^tp^, OAS3^tp^, OASL^tp^) were also overexpressed.

Regarding the IFN α/β pathway, our data highlighted elevated expression of type I interferon stimulated genes (ISGs)^26–28^ that mediate first-line antimicrobial defense by inhibiting viral fusion (IFITM1^tp^, IFITM3^tp^), replication (MX1^tp^, MX2^tp^, RSAD2^t^) and translation (IFIT1^t^, IFIT2^tp^, IFIT3^tp^, IFIT5^t^). The ubiquitin-like modifier ISG15^tp^, which interferes with viral protein function^29^, together with its modulator USP18^t^, was also upregulated.

Several IFN-γ induced genes also showed increased expression in our data, reflecting the activation of both innate and adaptive immune response. We observed overexpression of GBP proteins (GBP1^tp^, GBP3^t^, GBP4^tp^, GBP5^tp^, GBP6^tp^), which promote lysis of pathogen-containing vacuoles or bacteriolysis and inflammasome activation^30^, along with TRIM proteins (TRIM14^t^, TRIM21^t^, TRIM22^tp^), which target viral proteins acting as E3 ubiquitin ligases^31^. Other upregulated IFN-γ induced genes such as CIITA^t^, master regulator of MHC class II transcription^32^, FCGR1A^tp^, IFI30^p^ and ICAM1^tp^, enhance antigen presentation and the formation of the immunological synapse^33–35^.

In parallel, we observed enhanced expression of IL-4 and IL-13 signaling, indicative of Th2 cell activation (**Figure 1A, Supplementary Table S3**). Indeed, IL-13^t^, type I (IL4R^t^ and IL2RG^t^) and type II IL-4 receptor (IL4R^t^ and IL13RA1^p^), as well as the decoy receptor IL13RA2^t^ were increased. Upregulation of the upstream signal transducer JAK3^t^ further supports activation of this pathway^36^.

Our bulk transcriptomic dataset also revealed significant upregulation of other interleukins across distinct families in lesional samples. Members of the IL-1 family (IL-1β, IL-33, IL-36α, IL-36γ), the IL-17 family (IL-17A, IL-17C, IL-17F), and the IL-20 family (IL-19, IL-20, IL-24), as well as IL-6 and IL-23A, were all markedly induced, with IL-19 exhibiting the highest fold change (log₂FC = 9). Several CC (CCL2/4/5/8/17/20) and CXC (CXCL1/2/6/8/9/10/11) chemokines were also upregulated.

To localize upregulated pathways in HHD skin, we performed spatial RNA sequencing (Visium assays) on skin sections from 3 healthy donors and 2 HHD patients, profiling both lesional and non-lesional areas. After quality control, we detected between 3,740 (NL samples) and 5,282 (HC samples) spatially resolved spots with an average of 328 genes and 402 UMIs per spot. As shown in the spatial transcriptomic maps in **Figure 1D**, the IFN signaling pathway exhibited a significantly higher activity score in lesional skin compared with healthy and non-lesional skin, with preferential localization in the epidermal layers. We also mapped IFN signal transducers (JAK1, STAT1, STAT2), type I (ISG15, IFI6, IFI27) and type II IFN induced genes (HLA-DRA), confirming the increase in their expression in lesional skin *versus* healthy control (**Supplementary Figure S1**). By contrast, the IL-4/IL-13 signaling pathway score showed only a modest increase in lesional samples (**Supplementary Figure S2**).

### 2. scRNA-seq defines cell populations driving the IFN-response in HHD lesional skin

To dissect the cell-type composition of HHD lesions, we conducted a single-cell RNA sequencing (scRNA-seq) study of biopsies from lesional and non-lesional skin from 4 patients and 4 healthy individuals. We obtained a total of 53,760 cells (HC n= 17,193 cells, NL n=7,528 cells, L n=29,039 cells,) with an average of 2,970 genes and 11,722 transcripts detected per cell. Uniform Manifold Approximation and Projection (UMAP) identified a total of 19 main cell clusters, annotated using established canonical marker genes **(Figure 2A, Supplementary Figure S3)**. They included i) epithelial cells (basal keratinocytes, suprabasal keratinocytes, granular keratinocytes, melanocytes); ii) stromal cells (structural fibroblasts, inflammatory fibroblasts); iii) vascular cells (pericytes, lymphatic endothelial cells, vascular endothelial cells, smooth muscle cells); iv) immune cells (mast cells, natural killers cells, CD8+ T cells, CD4+ T cells, Langerhans cells, B cells, dendritic cells, monocytes and macrophages, plasmacytoid dendritic cells). Relative proportion of each cell type across the three conditions revealed a striking increase in immune cells in lesional skin, including innate (plasmacytoid dendritic cells, natural killer cells, dendritic cells) and adaptive immune populations (B cells, CD4+ T cells and CD8+ T cells) (**Figure 2B**). Notably, plasmacytoid dendritic cells (pDCs) were detected exclusively in lesional samples. In contrast, Langerhans cells appear to be reduced in the lesions. Compared to healthy controls, lesional skin also exhibited an increased number of inflammatory fibroblasts, pericytes, vascular endothelial cells and melanocytes, consistent with post-inflammatory hyperpigmentation. Moreover, the analysis highlighted an altered keratinocyte composition in both lesional and non-lesional skin.

**Figure 2.**
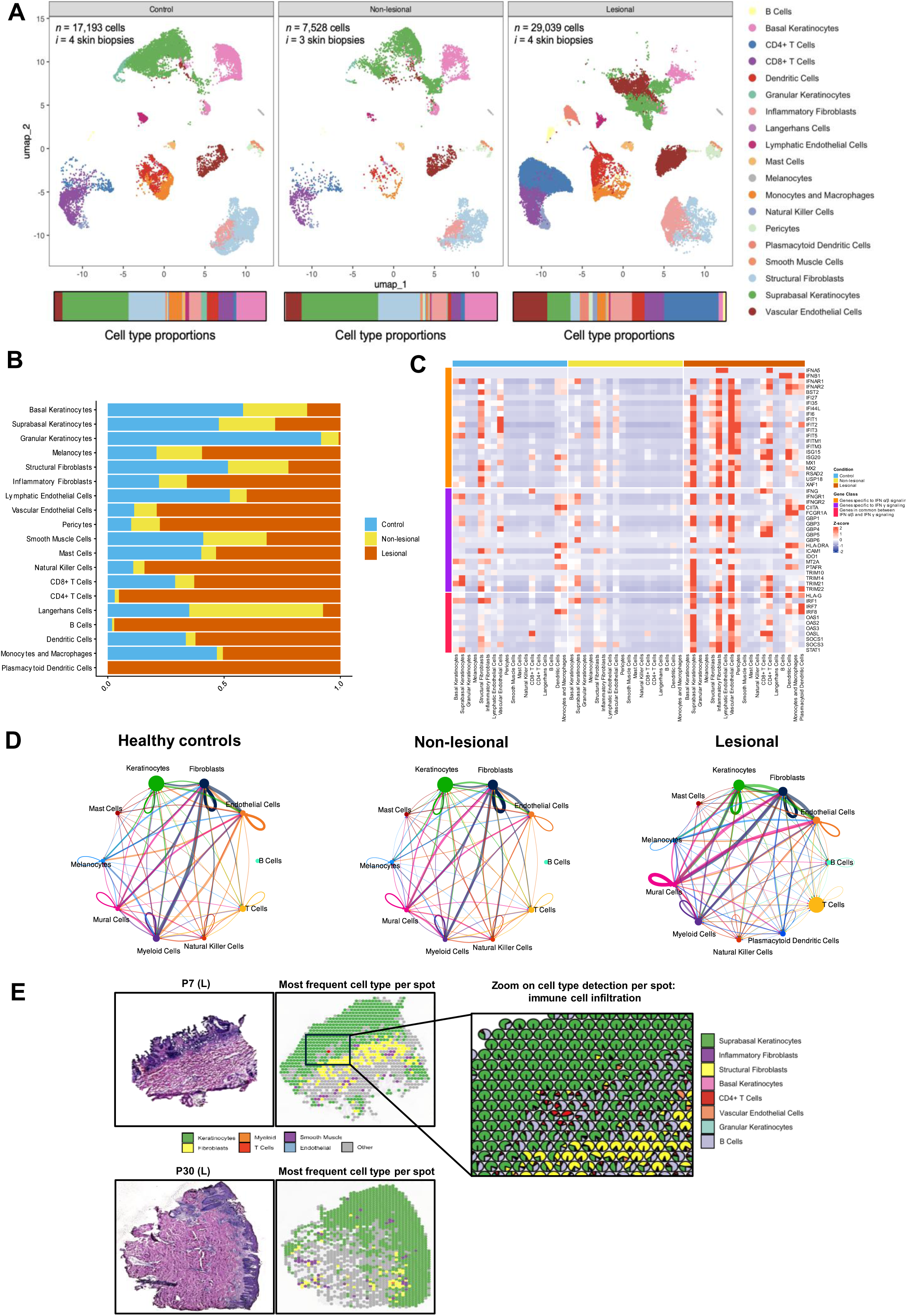
Cell populations associated with the IFN signature in HHD lesional skin. **(A)** UMAP projections of scRNA-seq data from control (*n* = 17,193 cells, 4 biopsies), non-lesional (*n* = 7,528 cells, 3 biopsies) and lesional skin (*n* = 29,039 cells, 4 biopsies). Bars below indicate relative proportions of each cell type across conditions. **(B)** Stacked barplots showing cell type composition stratified by condition (control, non-lesional, lesional). **(C)** Heatmap of type I and type II IFN signature gene expression across cell types in each condition. **(D)** Circle plots depict intercellular communication with each node representing a cell cluster. Node size indicates the total number of interactions in which the cluster participates, while edge thickness reflects interaction strength. **(E)** Visium SD assays of representative lesional skin sections. Left and down: H&E-stained tissue sections and spatial maps showing the most frequent predicted cell type per spot, revealing the spatial organization of keratinocytes, fibroblasts, T cells, myeloid cells, endothelial cells and smooth muscle cells. Right: higher-magnification view highlights immune cell infiltration and local heterogeneity in cell type composition within the epidermis in lesional skin.

To pinpoint the cellular drivers of the pronounced IFN response observed in HHD lesions, we measured the expression of IFN-related genes in each cell type population across the different conditions (HC, NL, L) (**Figure 2C**). In the lesional area, different cell types contributed distinctively to IFN signaling by overexpressing IFN molecules and/or interferon-stimulated genes (ISGs). Inflammatory fibroblasts, lymphatic endothelial cells, and CD4+ T cells expressed IFNA5. IFNB1 was produced by plasmacytoid dendritic cells, dendritic cells, and B cells. CD8+ T cells were the primary source of IFNG, with minor contribution from CD4+T cells and natural killer cells. Many of these populations act both as producers and responders to IFNs. Immune cells, including CD4⁺ and CD8⁺ T cells, dendritic cells, plasmacytoid dendritic cells, monocytes and macrophages, showed higher expression of type I and type II IFN-induced genes, confirming the well-established role of IFN as orchestrator and amplifier of innate and adaptive immunity^37^. Beyond immune cells, overexpression of ISGs was also observed in epithelial cells (suprabasal keratinocytes and, to a minor extent, basal keratinocytes and melanocytes), stromal cells (inflammatory fibroblasts) and vascular cells (pericytes and endothelial cells), underscoring the involvement of multiple cellular compartments in the IFN-coordinated inflammatory response. scRNA-seq based inference of ligand-receptor (L-R) interactions between cell clusters across skin states was performed to define cell-type specific crosstalk in lesional skin. Circle network plots, reported in **Figure 2D**, illustrate intercellular communication. Firstly, compared with healthy and non-lesional skin, lesional samples showed the inclusion of B cells in the communication network, together with pDCs, detected only in lesions. Moreover, T-cell node size was markedly increased, along with that of endothelial cells, highlighting their role as central hubs in lesional intercellular communication. In addition, as suggested by edge thickness, stronger interactions were observed among keratinocytes, vascular cells (mural and endothelial cells) and stromal cells (fibroblasts), pointing to an enhanced multicompartment network within the lesional tissue.

To define the spatial distribution of the main cellular drivers of the IFN response, we integrated scRNA-seq with Visium spatial transcriptomics by deconvolving each spot using reference cell type signatures. This analysis revealed a compartmentalized tissue architecture, with clear segregation of epidermal and dermal populations. Importantly, immune cells displayed non-random, spatially restricted enrichment patterns, forming localized infiltration zones rather than diffuse distribution across the tissue (**Figure 2E**). Regions at the dermal-epidermal interface showed increased cellular heterogeneity, suggesting areas of active immune-tissue interactions. Despite these insights, the limited resolution of Visium, which captures multiple cells simultaneously within the same spot, does not allow precise delineation of cellular neighborhoods and direct cell-cell interaction quantification, motivating the use of higher-resolution spatial transcriptomics.

### 3. High-resolution spatial transcriptomics maps IFN-driven inflammatory niches in lesional skin

To map the cellular landscape of HHD lesional skin at single-cell resolution, we performed Xenium spatial transcriptomics analysis. Cell types were assigned using gene expression signatures defined by scRNA-seq. As shown in **Figure 3A**, lesional *versus* non-lesional skin exhibited a striking enrichment of immune, stromal and vascular cells.

**Figure 3.**
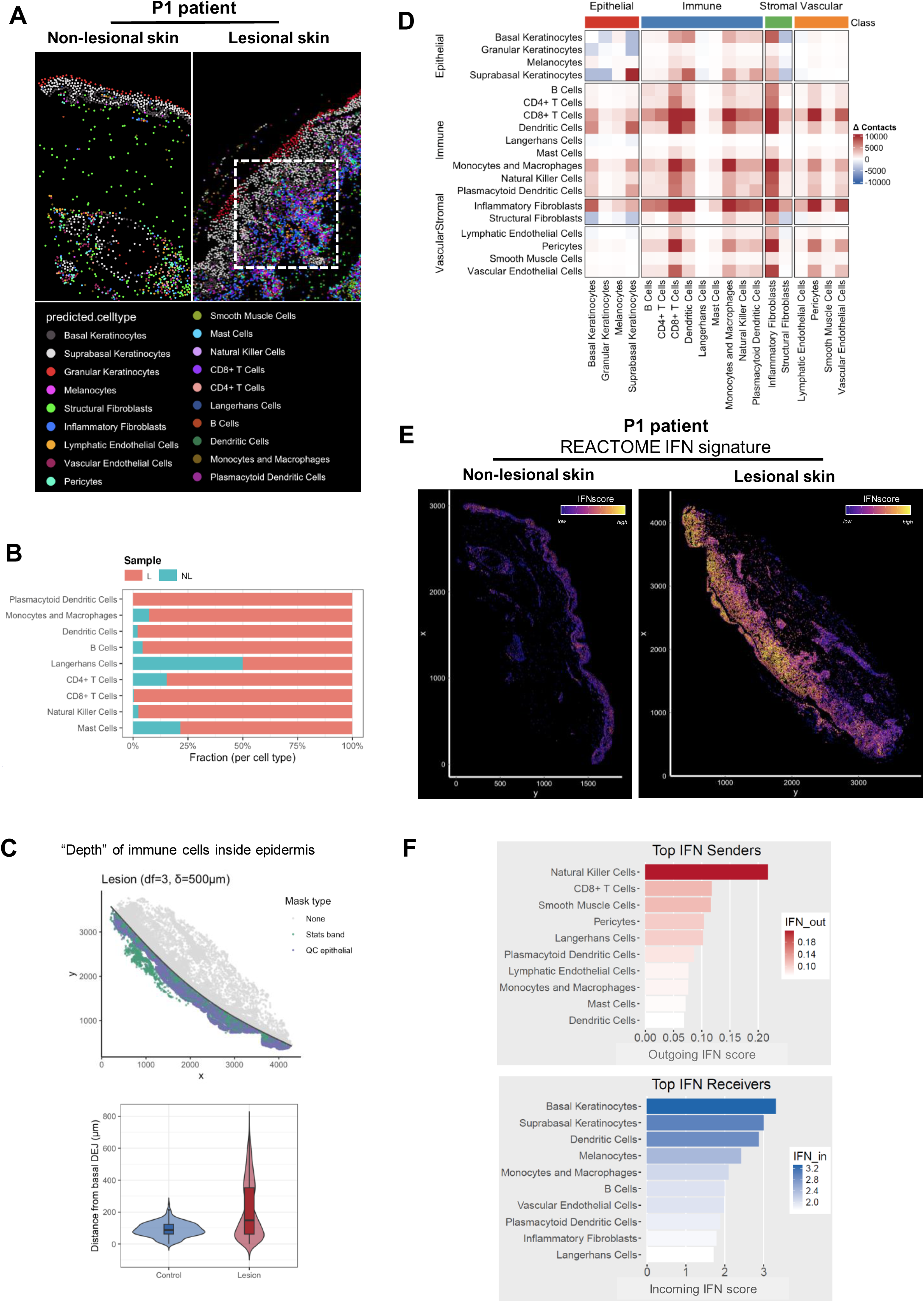
Single-cell resolution mapping of inflammatory niches in lesional skin. **(A)** Xenium assay for a patient biopsy, showing non-lesional (left) and lesional (right) skin. Cells are colored by predicted cell type, derived by label transfer from the matched scRNA-seq dataset. **(B)** Barplot of immune cell type proportions per condition (lesional vs non-lesional) for patient P1. **(C)** Quantification of the spatial depth of immune cells within the epidermis. Left: mask showing the epidermis-only selection, used to restrict all measurements to the epidermal compartment. The basal keratinocyte layer is defined as depth = 0 and serves as the reference boundary. Distances from this boundary to immune-cell centroids are computed to derive infiltration depth. Right: violin plot summarizing the distribution of immune-cell depth from the basal layer in control vs lesional epidermis, demonstrating deeper infiltration in lesions. **(D)** Differential cell-cell proximity analysis between lesional and non-lesional skin. Heatmap shows the change in adjacency frequency (Δ L/NL) across major cell type pairs, stratified by epithelial, immune, stromal and vascular classes. **(E)** Spatial transcriptomics maps (Xenium assays) showing IFN signaling pathway activity scores in representative patient non-lesional and lesional skin sections. **(F)** Bar plots illustrate outgoing and incoming IFN signaling activity score by cell type.

In line with the scRNA-seq data described above, cell-type proportion analysis across conditions revealed a pronounced increase of almost all immune cell subtypes in lesional samples, including myeloid populations (plasmacytoid dendritic cells, monocytes, macrophages, dendritic cells and mast cells), natural killer cells, T cells (CD4+ and CD8+) and B cells (**Figure 3B**).

Accordingly, immune cells expanded beneath and within the epidermal layers of the lesional skin (**Supplementary Figure S4**). To quantify positioning relative to the epidermis, we defined the dermal-epidermal junction (DEJ) from basal keratinocyte coordinates and fitted a smooth reference curve. Immune cells lying above the predicted DEJ were classified as epidermal; we computed the vertical distance between each immune cell centroid and the curve as a measure of infiltration into the epidermal compartment. Lesional skin showed greater epidermal immune infiltration than non-lesional skin (**Figure 3C**).

To map local cellular neighborhoods within the lesional tissue and to define the cellular composition of the infiltrate, we performed differential cell-cell proximity analysis in L *versus* NL skin (**Figure 3D**). First, this highlighted hallmark features of acantholysis in lesional samples, with reduced contacts between suprabasal-basal and suprabasal–granular keratinocytes. Notably, contacts among suprabasal keratinocytes themselves were increased, consistent with non-acantholytic, hyperplastic epidermal areas. Suprabasal keratinocytes, basal keratinocytes, and to a lesser extent, melanocytes showed augmented proximity with CD8+T cells, dendritic cells, monocytes, macrophages, natural killer cells and pDCs. Higher adjacence between these epithelial cells and inflammatory fibroblasts was also observed. The aforementioned immune cells, together with B cells and CD4+ T cells, displayed increased proximity with inflammatory fibroblasts, pericytes, and vascular endothelial cells, delineating the dermal immune infiltrate. Spatial adjacence between the same stromal and vascular populations was also enhanced.

Xenium analysis confirmed a markedly higher IFN signaling pathway activity score in lesional skin compared with non-lesional sample (**Figure 3E**).

To assess functional cell-cell communication, IFN-specific ligand-receptor interactions were evaluated across cell populations in L *versus* NL skin. (**Figure 3F**). While NKs, CD8+ T cells and pDCs were confirmed as IFN producers, additional cell types were also identified, including smooth muscle cells, pericytes and Langerhans cells. Beyond immune cells, epithelial cells (basal and suprabasal keratinocytes, melanocytes) were the top IFN receivers, together with vascular endothelial cells, although to a lesser extent. These results, obtained at the spatial level, globally support and complement scRNA-seq results from dissociated cells.

### 4. Ruxolitinib treatment resolved HHD lesions

To translate our findings into a therapeutic framework, we applied immune-related modules (type I IFN, type II IFN, eosinophilic, macrophagic, neutrophilic, Th1, Th2 and Th17) (**Supplementary Table S4**) established by Seremet, T. *et al.*^38^ and Ayers, M. *et al.* ^39^ to our bulk transcriptomic dataset, enabling precise delineation of immune pathway activity across our patient cohort. IFN signature emerged as the prominent axis, with concurrent activation of macrophagic, neutrophilic, Th1, and Th17 modules, whereas eosinophilic and Th2 signals were heterogeneous, highlighting a complex inflammatory network (**Supplementary Figure S5A**). To explore therapeutic candidates, we queried significantly dysregulated immune module genes against the Open Targets Platform ^40^ to identify molecular interaction partners and generate a comprehensive target-drug map. The search was restricted to Food and Drug Administration approved drugs to facilitate rapid clinical repositioning. As a result, 177 approved drug candidates were identified and ranked based on their total number of targets (**Supplementary Figure S5B, Supplementary Table S5**). Given the co-expression of multiple inflammatory pathways, we implemented three criteria for drug selection: (1) potent inhibition of IFN signaling, (2) broad coverage of secondary dysregulated modules, and (3) suitability for topical delivery to maximize local efficacy while minimizing systemic exposure. Based on this approach, we identified ruxolitinib as the optimal therapeutic candidate. Notably, ruxolitinib was the only agent among the top seven ranked groups available in a topical formulation, directly addressing our prioritization of local delivery. As a JAK1/2 inhibitor, ruxolitinib allows for the targeting of the predominant IFNs axes and the secondary immune modules activated in lesional samples. Accordingly, we treated ten HHD patients with active lesions resistant to standard treatments using off-label topical ruxolitinib (**Supplementary Table S6**).

Clinical efficacy was evaluated through clinical progression of the lesions and patient-reported outcomes (PROs). In nine patients, the effect was rapid as early as the first 7 days, showing substantial improvement in pain and oozing, resolution of vesicopustules, erosions and fissures, with re-epithelialization within a month (**Figure 4A**). One patient (HH4) with a secondary bacterial skin infection showed partial response, even after antibiotic treatment. Treatment was well-tolerated, although patient HH8 developed a local herpes infection confirmed by PCR, who responded well to oral valaciclovir and topical acyclovir. Responses remained durable over up to nine months of follow-up, persisting after de-escalation from twice-daily to once-weekly dosing with eventual discontinuation of treatment.

**Figure 4.**
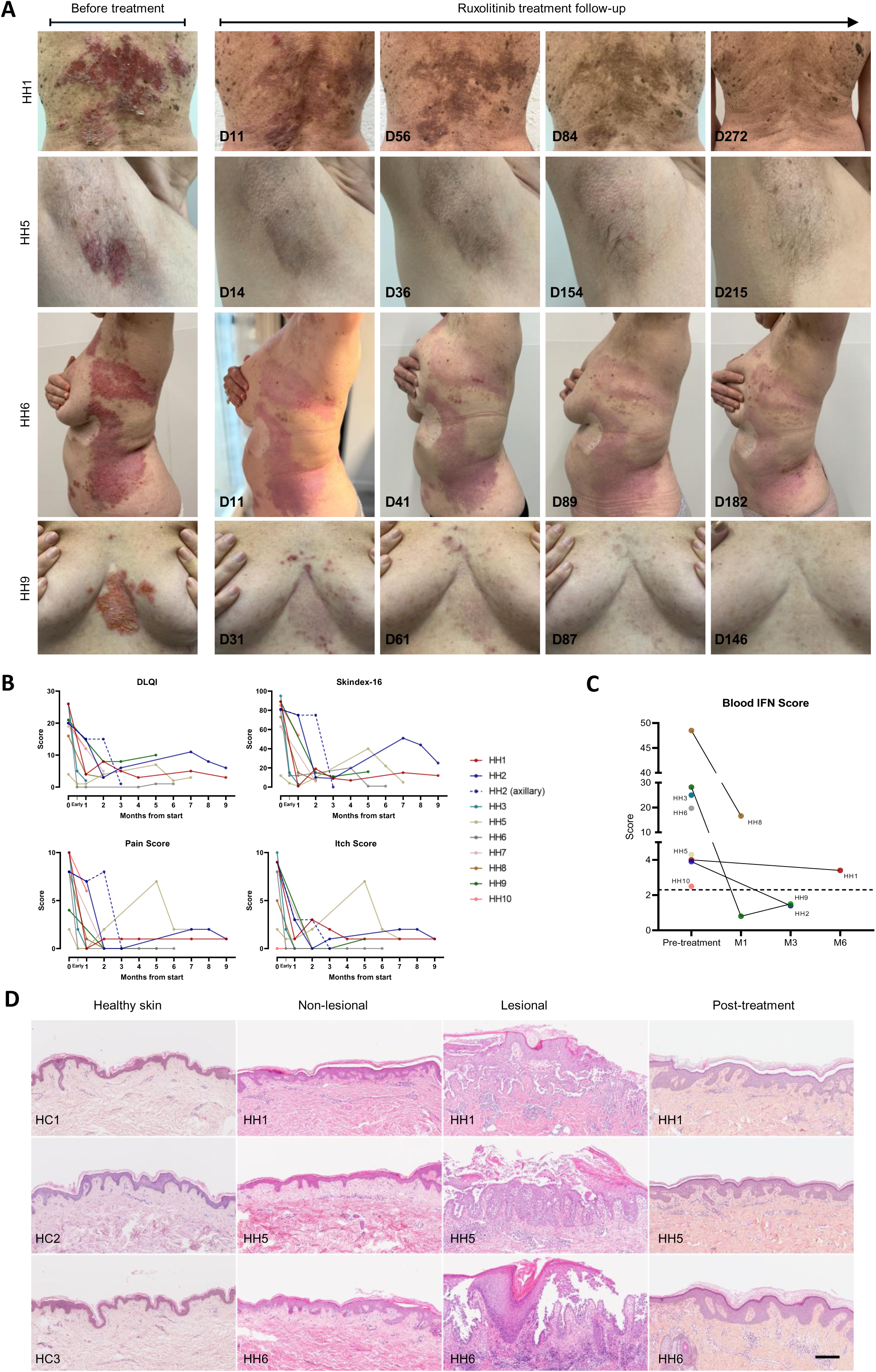
Clinical and histopathological improvement of Hailey-Hailey (HH) lesional skin following ruxolitinib treatment. **(A)** Representative clinical photographs of HH patients before treatment and during follow-up after ruxolitinib treatment initiation. **(B)** Patient-reported outcomes, including Dermatology Life Quality Index (DLQI), Skindex-16, pain, and itch scores, assessed before treatment and during follow-up after treatment initiation (early: days 1-14; months 1-9). **(C)** Blood type I interferon scores measured in peripheral blood based on transcript levels of a targeted panel of interferon-stimulated genes. Individual data points correspond to specific HH patients (pre-treatment: n=8; months 1, 3 and 6 after treatment initiation: n=2, 2 and 1, respectively). Black lines connect values from the same patient. The dashed line indicates the positive IFN signature threshold score of 2.3. **(D)** Representative haematoxylin and eosin (H&E) staining of skin sections from healthy controls and HH patients. Post ruxolitinib-treated lesional skin from HH1, HH5, and HH6 was biopsied on days 41, 57, and 26 after treatment initiation, respectively. Scale bar = 200 µm.

Dermatology Life Quality Index (DLQI), Skindex-16, Numeric Rating Scale (NRS) for pain and itch were used to assess subjective clinical outcomes and quality of life (QoL) (**Figure 4B**). Patients reported a rapid decrease in median DLQI (from 20 [4-26] to 4 [0-15]) and Skindex-16 scores (from 81 [12-95] to 12 [1-75]) within 1-2 months of treatment. Clinical improvement was sustained, with DLQI remaining low with a median of 2 [1-10] at month 5-6, consistent with small or no impact on QoL. Patients described reduced lesion-related emotional stress, associated with a decrease in the occurrence of new lesions.

In parallel, symptom trajectories mirrored the QoL improvements (**Figure 4B**). By 1-2 months, patients showed a drastic reduction in median NRS pain (from 8 [2-10] to 0 [0-8]) and itch scores (from 8.5 [0-10] to 0 [0-3]), which remained low at 6-9 months (1.5 [0-2] for pain and 1 [0-2] for itch).

Strikingly, very early responses were observed in HH3 and HH6, who showed notable improvements in PROs within the first post-treatment week, that were sustained across follow-up periods. Patients HH1, HH2, and HH5 similarly showed a rapid response to ruxolitinib treatment of the initial lesions and new active lesions developed during follow-up outside the initially treated areas (**Supplementary Figures S6, S7 and S8B**). This improvement was accompanied by a decrease in HH1, HH2 and HH5 scores, after prior increment at month 2, 7, and 5, respectively (**Figure 4B**). In contrast, in HH2, the partial improvement of an axillary region after 1 month of ruxolitinib treatment was associated with bacterial infection (**Supplementary Figure S8A**). Resuming treatment with ruxolitinib, after local antibiotic treatment (mupirocin), led to marked clinical improvement, with DLQI decreasing to 1 by month 3.

As demonstrated by our omics data, IFN signaling is a key driver of cutaneous inflammation in patients. We therefore assessed whether HHD is also associated with systemic IFN-responsive signature and whether this is modulated by ruxolitinib treatment. To address this, we measured a blood IFN score based on transcript levels of a targeted panel of type I interferon-stimulated genes (*IFIT1, ISG15, RSAD2, IFI27, SIGLEC1* and *IFI44L*). Before treatment, patients showed a positive but heterogeneous IFN signature, with a median score of 12 [2.5–48.5] (**Figure 4C**), consistent with systemic IFN-associated response. Notably, after initiation of topical ruxolitinib, IFN scores declined overall, supporting a role for the cutaneous lesions as a primary driver of inflammation.

To document the histopathological remodeling in clinically healed areas following ruxolitinib treatment, we compared healthy control, non-lesional, baseline lesional and post-treatment (PT) skin samples (**Figure 4D and Supplementary Figure S9**). While HC and NL skin showed normal stratification and minimal dermal immune cells, baseline lesional sections displayed the hallmarks of Hailey-Hailey disease, with epidermal acantholysis accompanied by epidermal thickening and a dense inflammatory cell infiltrate. Strikingly, post-treatment samples showed clear reduction in inflammatory cell infiltrates with restoration of cell-to-cell adhesion and decrease in epidermal thickness. Together, these histological findings, which parallel the clinical improvement observed after topical ruxolitinib, demonstrate that clinical healing is accompanied by restoration of epidermal architecture and attenuation of cutaneous inflammation.

### 5. Ruxolitinib abolished IFN response, reduced immune cell infiltration and rescued epidermal architecture

Immunofluorescence staining for phosphorylated STAT1 (pSTAT1) and phosphorylated STAT3 (pSTAT3), confirmed engagement of IFN downstream JAK-STAT signaling in HHD lesional samples, with minimal to no signal in healthy control and non-lesional skin (**Figure 5A and B**). IRF9 and IDO1, which are interferon-responsive markers associated with type I ^41^ and type II interferon signaling^42^ respectively, were highly expressed in HHD lesions, with IRF9 predominantly localized to keratinocytes and IDO1 to dermal cells (**Figure 5C and D**) in HHD lesions. Following topical ruxolitinib treatment, pSTAT1, pSTAT3, IRF9 and IDO1 signals returned toward baseline in post-treatment skin, indicating resolution of the interferon inflammatory response after JAK inhibition.

**Figure 5.**
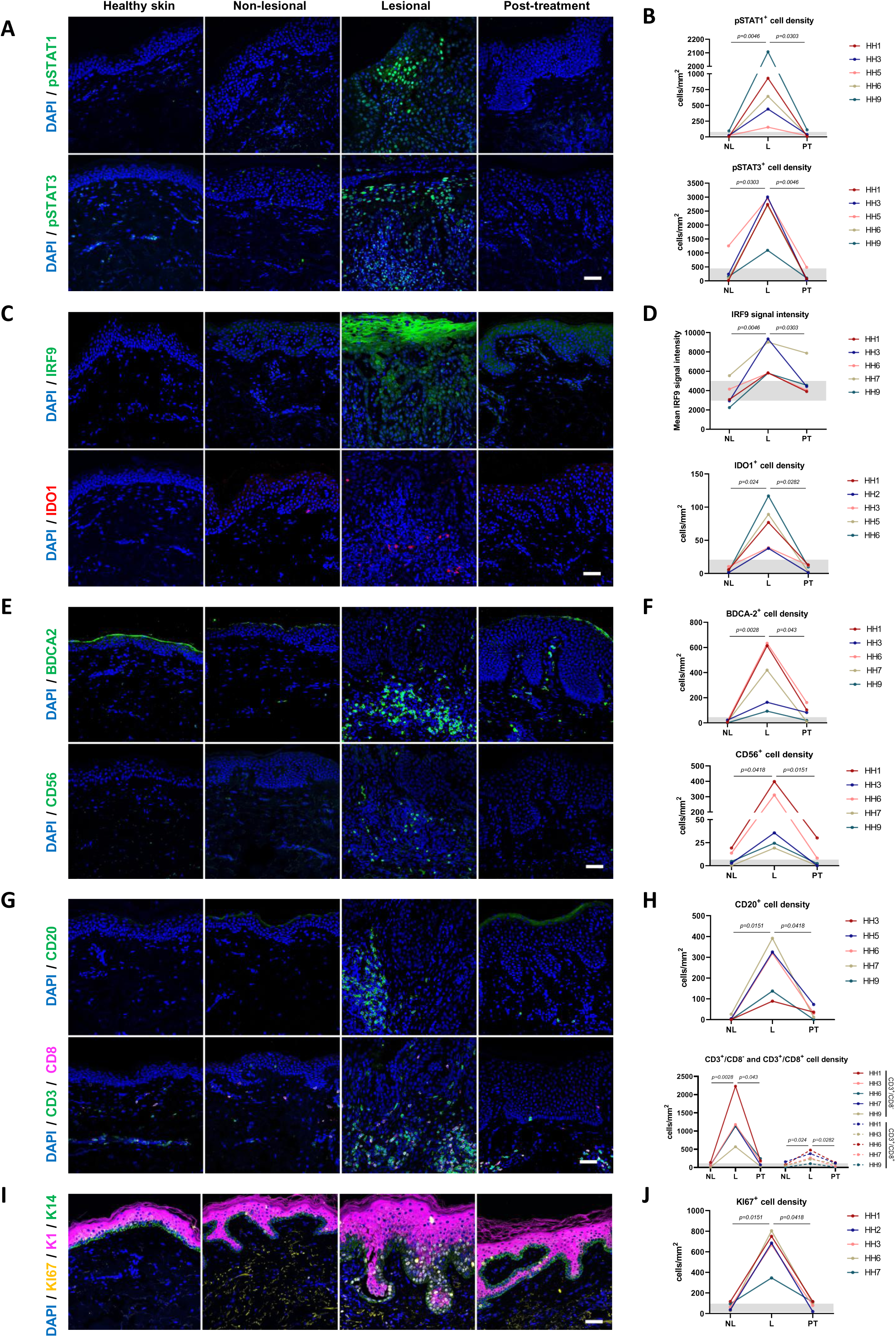
Ruxolitinib abolishes IFN response, attenuates immune infiltration and restores epidermal differentiation and proliferation in Hailey-Hailey disease. **(A)** Representative immunofluorescence images of phosphorylated STAT1 (upper panel, pSTAT1, green) and phosphorylated STAT3 (lower panel, pSTAT3, green) with **(B)** corresponding quantification of positive cell densities (cells per mm^2^) across patient-matched non-lesional (NL), lesional (L), and post-treatment (PT; post-ruxolitinib treatment) skin. **(C)** Representative immunofluorescence images of type I interferon-inducible marker IRF9 (upper panel, green) and type II interferon-inducible marker IDO1 (lower panel, red), with **(D)** corresponding quantification of IRF9 mean signal intensity and IDO1 cell density (cells per mm^2^) across patient-matched NL, L and PT skin. Representative immunofluorescence images of **(E)** innate immune cell populations, including plasmacytoid dendritic cells (upper panel, BDCA2, green) and natural killer cells (lower panel, CD56, green); and **(G)** adaptive immune cell populations showing B cells (upper panel, CD20, green) and T-cell subsets (lower panel, CD3, green; CD8, magenta). Respective quantification of **(F)** positive innate immune cells and **(H)** positive adaptive immune cells densities (cells per mm^2^) across patient-matched NL, L and PT skin. **(I)** Keratinocyte proliferation (Ki67, yellow) with **(J)** associated quantification of KI67 positive cell density (cells per mm^2^) in NL, L, and PT skins. Gray shaded areas represent the range measured in healthy controls (n=3). Nuclei were counterstained with DAPI (blue). Scale bars = 50 µm. Data points represent individual patients (n=5). Differences between groups were evaluated using the Friedman test, followed by multiple-comparison adjustment using the two-stage linear step-up procedure of Benjamini, Krieger and Yekutieli. Statistical significance was defined as p < 0.05.

Consistently, immunostaining also showed that ruxolitinib attenuates immune infiltration in HHD lesions. Before treatment, lesional skin displayed marked accumulation of both innate and adaptive immune cells. Regarding innate populations, dense infiltrates of IFN-producing pDCs (BDCA2⁺) and NK cells (CD56⁺) were detected (**Figure 5E and 5F**), which significantly reduced after treatment; residual pDCs persisted. In addition, a prominent infiltration of monocytes (CD14⁺), macrophages (CD68⁺), and neutrophils (MPO⁺) was observed in lesional skin **(Supplementary Figure S10A-D**). Following ruxolitinib treatment, residual CD14⁺ and CD68⁺ cells remained detectable, whereas neutrophils were largely absent. Notably, Langerhans cells (CD207⁺) were depleted in the lesional epidermis, suggesting inflammation-driven migration toward skin-draining lymph nodes^43^, and repopulated after treatment. The adaptative compartment was also expanded in lesional samples, with accumulation of B (CD20⁺) and T cells (**Figure 5G and 5H**); including a relative enrichment of CD3⁺CD8⁻ (presumably CD4⁺ helper) over CD3⁺CD8⁺ cytotoxic T cells. B and T cell populations were significantly reduced after ruxolitinib treatment, with a small residual fraction persisting.

Given that the post-treatment samples were obtained at a median of 35 [26-57] days after treatment initiation, these findings are consistent with ongoing resolution of the inflammatory infiltrate.

We also observed restoration of normal epidermal proliferation, differentiation, and barrier integrity following ruxolitinib treatment. In the healthy epidermis, keratinocytes follow a tightly regulated differentiation program and switch from basal keratins (K5/K14) to suprabasal keratins (K1/K10) as they migrate upward. In HHD lesional skin, the first suprabasal layers expressed K14 in several regions instead of K1, indicating impaired early-stage differentiation (**Figure 5I**). Concurrently, lesional keratinocytes were hyperproliferative, as shown by a significant increase in KI67+ cells in basal and suprabasal layers (**Figure 5I and 5J**). In addition, expression of late-stage differentiation, barrier-associated markers filaggrin (FLG) and loricrin (LOR) were also disrupted. While FLG staining appeared weak and discontinuous in the granular layer, LOR expression was aberrant, extending beyond the granular layer (**Supplementary Figure S10E**). In post-treatment skin, we observed substantial normalization of these features: K14 became confined to the basal layer; KI67+ cells decreased in number and were restricted to basal-layer levels; FLG and LOR staining were restored in continuous, linear patterns, comparable to healthy control and non-lesional skin.

## DISCUSSION

All omics layers converged on a striking IFN signature in HHD lesions, highlighting its role as a central driver of the inflammatory process. Integration of single-cell and spatial transcriptomics revealed the presence of inflammatory niches, where immune, epithelial, vascular, and stromal cells are engaged in a dynamic crosstalk to amplify the IFN-driven immune response.

**Figure 6 (upper panel)** depicts the proposed inflammatory mechanism underlying Hailey-Hailey disease.

**Figure 6.**
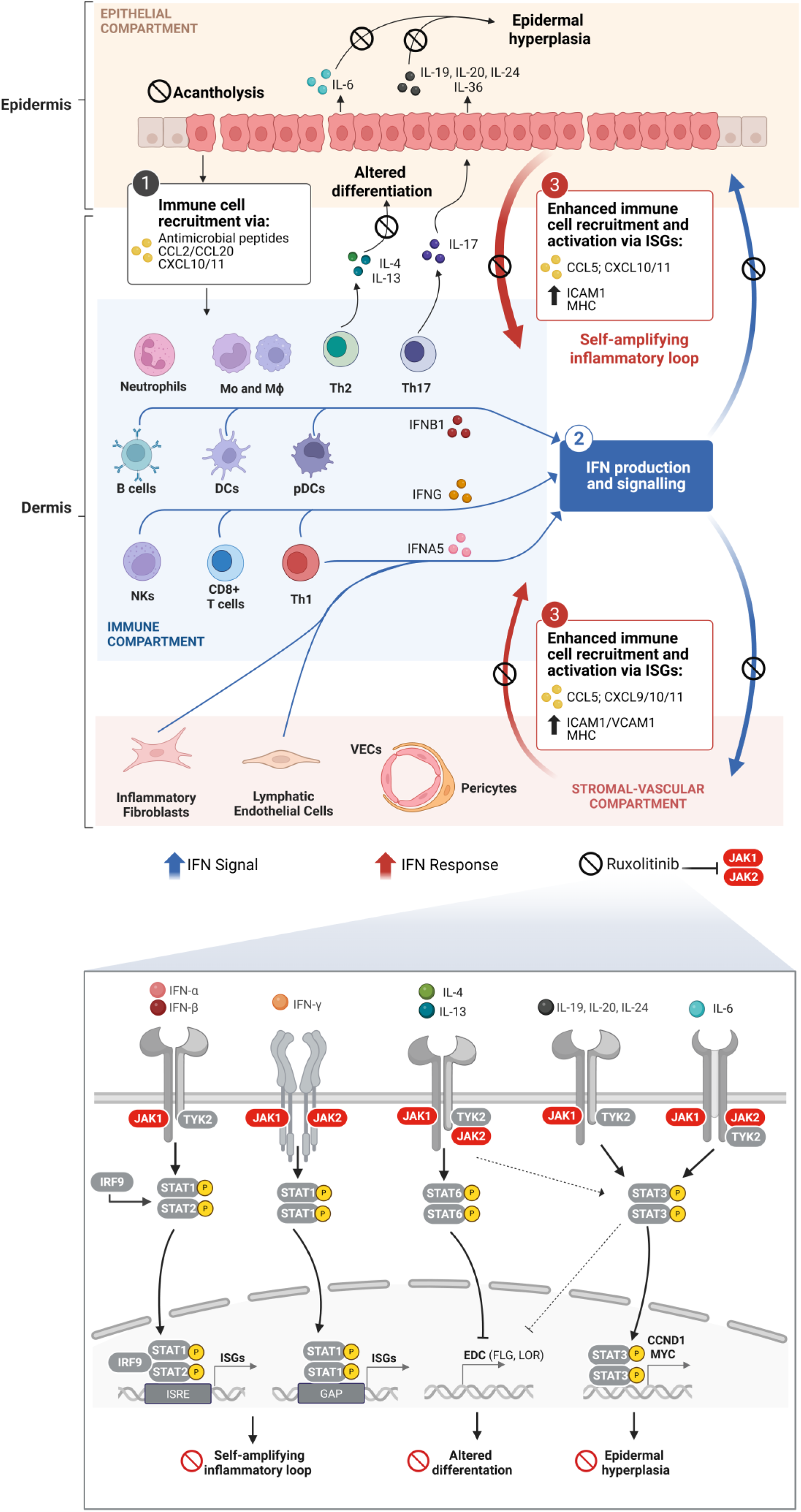
Proposed inflammatory mechanism underlying Hailey-Hailey disease and its modulation by ruxolitinib. (1) Keratinocytes in acantholytic lesional skin could initiate immune cell recruitment via antimicrobial peptides, interleukins and chemokines (CCL2/20, CXCL10/11). (2) Among recruited immune cells, plasmacytoid dendritic cells (pDCs), dendritic cells (DCs), and B cells produce type I interferons, whereas CD8⁺ T cells and natural killer (NK) cells produce type II interferon, with CD4⁺ T cells contributing to both. (3) Keratinocytes, together with vascular cells and inflammatory fibroblasts, respond to interferon signals by amplifying chemokine production (CCL5, CXCL9/10/11) and upregulating MHC and adhesion molecules, thereby enhancing immune cell recruitment and establishing a self-amplifying inflammatory loop. IL-6 together with IL-17A-induced IL-20 family cytokines and IL-36 contribute to epidermal hyperplasia, whereas IL-4 and IL-13 impair epidermal differentiation. Ruxolitinib, by targeting locally JAK1 and JAK2, inhibits interferon signaling and the self-amplifying inflammatory loop, restores cell-cell adhesion, normal keratinocyte differentiation and proliferation.

Acantholytic keratinocytes could act as initiators of immune cell recruitment, as suggested in our dataset by the upregulation of antimicrobial peptides, including cathelicidin (CAMP) and psoriasin (S100A7), alongside a broad array of interleukins and chemokines. This repertoire promotes the recruitment of monocytes and dendritic cells via CCL2; elevated levels of CXCL10 and CXCL11 enhance trafficking of Th1 cells, CD8⁺ T cells, and NK cells, whereas CCL20 facilitates the recruitment of Th17 cells^44–47^. IL-33 is also upregulated in our data and promotes Th2-type cytokine-associated immune responses^48^.

The immune compartment was found to be the principal source of IFN signaling. In line with the literature, plasmacytoid and conventional dendritic cells contributed to type I interferon production^49^, whereas CD8⁺ T cells, NK cells, and CD4⁺ T cells acted as sources of IFN-γ^22^. Unexpectedly, we also detected type I IFN expression in B cells and CD4⁺ T cells, previously reported only in limited pathological contexts^50,51^. Consistent with previous studies^52–54^, inflammatory fibroblasts and vascular cells were also found to produce type I IFN.

Within the lesional microenvironment, immune cells acted not only as producers but also as responders to IFN, highlighting interferon as a pivotal amplifier of innate and adaptive immunity. Indeed, type I and type II IFNs are known to promote antigen-presenting cell (APC) maturation, enhance NK cell cytotoxicity, drive Th1 polarization of CD4⁺ T cells, and increase the proliferation and effector functions of CD8⁺ T cells ^37,55^.

Beyond immune cells, epithelial cells (basal and suprabasal keratinocytes, melanocytes) were the primary recipients of IFN signaling, with vascular cells and inflammatory fibroblasts also responding, although to a lesser degree. Upon IFN stimulation, these populations upregulate MHC molecules, adhesion molecules (ICAM-1 broadly, VCAM-1 in vascular and stromal cells), and chemokines (CCL5, CXCL9/10/11), driving further immune cell recruitment and amplifying local inflammation^56–58,52,59^. Thereby, collectively, our data support the existence of a multi-compartment IFN-driven signaling network that establishes a self-sustaining feed-forward amplification loop, underlying the relapse and chronicity of inflammation in HHD skin lesions. Based on these findings, we targeted the IFN–JAK–STAT axis and effectively treated nine HHD patients with the topical JAK1/2 inhibitor ruxolitinib.

We selected a JAK1/2 inhibitor based on its capacity to concurrently inhibit type I and II IFN signaling, as well as IL-4/IL-13, IL-6, and IL-20 cytokine pathways^60,61^, all of which are upregulated in lesions and likely to contribute to epidermal hyperplasia and barrier disruption seen in HHD (**Figure 6, lower panel**).

Ruxolitinib, which has already been approved for vitiligo^62^ and atopic dermatitis^63^, was preferred to oral JAK1/2 inhibitors due to minimal systemic exposure and a reduced risk of adverse events, including infections, musculoskeletal and connective tissue disorders, venous thromboembolism and neoplasms^64,65^.

Ruxolitinib effectively disrupted the IFN-driven inflammatory amplification loop, leading to a marked reduction in immune cell infiltration and restoration of keratinocyte homeostasis, including cell-cell adhesion, differentiation and proliferation. Resolution of inflammation may alleviate cellular stress, including ER and Golgi apparatus stress^66–69^, which has been implicated in acantholysis in HHD. In addition, the restoration of cell–cell adhesion may be partly mediated through STAT1 and STAT3 inhibition, which has been associated with increased expression of desmosome components and adherens junction proteins ^70,71^.

Reestablishment of intracellular junctions via cadherins is likely to improve early keratinocyte differentiation^72^. Inhibition of IL-4/IL-13 and IL-24 may enhance late keratinization through upregulation of EDC molecules (filaggrin and loricrin) via STAT3-and STAT6-dependent mechanisms^61^.

Suppression of STAT3 activation downstream of IL-6 and IL-20 family cytokines may also contribute to the normalization of keratinocyte proliferation through downregulation of proliferation-associated gene programs (CCND1, MYC)^73,74^.

The effectiveness of local JAK1/2 inhibition, coupled with the observed reduction of the systemic IFN signature, suggests that the cutaneous microenvironment acts as the primary driver of inflammation. The IFN signature in peripheral blood likely reflects the response of circulating immune cells to IFN produced locally and released into the systemic circulation, as reported in viral lung infections ^75^.

Of the ten treated patients, who were refractory to classical treatment, nine patients exhibited a remarkable positive response to ruxolitinib. Treatment was well-tolerated, and responses remained durable over up to nine months of follow-up, leading to a significant improvement in quality of life.

Of note, in case of suspected bacterial infections, which are frequent in HHD, especially in intertriginous regions (axillary, submammary and inguinal folds), topical and/or oral antibiotic therapy should be considered prior to ruxolitinib treatment.

Some patients developed new lesions at the periphery of affected areas treated with ruxolitinib, suggesting that perilesional skin is primed for inflammation. We concluded that topical applications should extend beyond the visible lesion to prevent peripheral expansion. These peripheral lesions, as well as new lesions distant from the initially treated areas, responded well to ruxolitinib treatment with rapid improvement, reduction in pain, and complete healing. Patients generally expressed a strong preference for repeated topical application rather than systemic therapy. However, in cases of multiple and/or extensive erosive and painful lesions, oral JAK inhibitors may be considered in the absence of contraindications, by closely monitoring potential side effects.

Our study provides a successful example of how a multiomic approach can directly lead to mechanism-based therapeutic strategies. Given the sustained efficacy in treated patients, our results highlight an effective and well-tolerated treatment for severe HHD. We believe topical ruxolitinib is a transformative treatment which could become the first line standard of care for Hailey-Hailey disease.

## METHODS

### Patient and healthy control biopsies

Collection of samples from healthy donors and patients with HHD was conducted in accordance with the Declaration of Helsinki and was reviewed and approved by the Internal Review Board (IRB) of the Imagine Institute for Genetic diseases (Paris, France). Patients were seen by A.H. at the Department of Genomic Medicine for Rare Diseases at The Imagine Institute for Genetic Diseases. The diagnosis of HHD was based on the clinical presentation and was confirmed by histopathological findings **(Figure 4D, Supplementary Figure S9 and S11)** and genetic identification of a heterozygous *ATP2C1* pathogenic variant. All patients and healthy controls were informed and provided their written consent for use of samples for research purposes including transcriptomic and proteomic assays. Skin tissue biopsies were obtained from 32 adult patients with HHD and 28 age-and sex-matched healthy controls (**Supplementary Table S1**). All samples were deidentifed. The samples are part of a registered Biobank at INSERM (INSERM DC-2019-3504).

### Patient treatment

We treated ten HHD patients with increased IFN signature at the transcriptomic and proteomic levels, and severe active clinical manifestations resistant to classical treatments (**Supplementary Table S6**). Off-label therapeutic regimen using a topical ruxolitinib 1.5% cream (Opzelura®, Incyte Corporation) was applied to affected areas. Treatment was started with twice-daily application until lesion resolution and then reduced to once daily for 1 week. Patients were off systemic treatment during the study.

### Bulk-transcriptomics analysis

5 mm full-thickness skin biopsies were collected and immediately submerged in RNAprotect Tissue Reagent (Qiagen, ref: 76106), incubated for 24 h at 4°C, and then stored at-80°C. Frozen samples were cryogenically pulverized using cell crusher kit (CellCrusher, ref: 538005) then immediately lysed in 1% β-mercaptoethanol/RLT buffer. Total RNA was extracted using RNeasy Fibrous Tissue Mini Kit (Qiagen, ref: 74704) according to the manufacturer’s instructions. RNA concentration and purity were assessed using the Expose spectrophotometer (Trinean). 400 ng of total RNA was used for library construction using the Universal Plus mRNA-Seq Kit (Tecan). Sequencing was performed on Illumina NovaSeq 6000 using S4 flow cell with a 100-10-10-100 cycle configuration (paired end: R1 = 100 bp, i7 = 10 bp, i5 = 10 bp, R2 = 100 bp). Bulk transcriptomic data were processed in R (v4.4.2) using RStudio (v2024.09.0). Differential expression analysis was conducted using the DESeq2 package (v1.46.0). Statistical significance was evaluated using Wald tests, followed by Benjamini-Hochberg correction.

### Proteomics analysis

#### Sample preparation

5 mm full-thickness snap-frozen skin biopsies were collected from lesional and non-lesional areas of 24 Hailey-Hailey patients. They were processed using BeatBox Tissue Kit 24x (PreOmics), following the addition of the lysis buffer (8 M urea, 5% SDS, 50 mM TEAB). After heating at 95° for 5 minutes and centrifugation at 14’000g for 10 minutes, protein dosage was performed using DC Protein Assay Kit (Bio-rad). S-Trap™ 96-well plate (Protifi, Hutington, USA) digestion was performed on 30µg of cell lysates according to manufacturer’s instructions. After elution, peptides were vacuum dried and resuspended in 75µl of 2% ACN, 0.1% formic acid in HPLC-grade water.

#### nanoLC-MS/MS analysis

400 ng were injected on a nanoElute2 (Bruker Daltonics, Germany) HPLC (high-performance liquid chromatography) system coupled to a timsTOF Pro2 (Bruker Daltonics, Germany) mass spectrometer. HPLC separation (Solvent A: 0.1% formic acid in water; Solvent B: 0.1% formic acid in acetonitrile) was carried out at 250nL/min using a packed emitter column (C18, 25 cm×75μm 1.6μm) (Ion Optics, Australia) and a 40min gradient elution (2 to 11% solvent B during 19min; 11 to 16% during 7min; 16% to 25% during 4min; 25% to 80% for 3min and finally 80% for 7min to wash the column). Mass-spectrometric data were acquired using the parallel accumulation of serial fragmentation (PASEF) acquisition method in DIA mode. diaPASEF settings were: mass range from 475 to 1000 m/z, mobility ranges from 0.85 to 1.27 V s/cm2 1/k0, cycle time estimate of 0.95s, mass width of 25Da and 21 mass steps for cycle.

#### Proteomics data analysis

Data analysis was performed using Spectronaut software (version 20.1). A search against the Homo sapiens (UniProtKB/Swiss-Prot) database (downloaded on 12 February 2025, 20417 entries) was performed using the directDIA+ (Deep) workflow with default settings. Bioinformatic analysis was performed using R (v4.4.0) and Rstudio (RStudio 2025.09.0). For statistical comparison, raw Label Free Quantification (LFQ) intensities were log2-transformed, and we set two groups corresponding to the experimental conditions (NL for Non-Lesional and L for Lesional), each containing 24 biological replicates. We then filtered the data to keep only proteins with valid values in at least 50% of the replicates in at least one experimental group. Next, missing values were imputed to fill missing data points by creating a Gaussian distribution of random numbers with a standard deviation of 33% relative to the standard deviation of the measured values and 1.8 standard deviation downshift of the mean to simulate the distribution of low signal values. Finally, limma-based statistics were performed between L and NL groups with logFC threshold set a-1 and 1 and Benjamini-Hochberg multiple test correction was applied at 5% to control false discovery rate.

### Integration of bulk RNA-seq and proteomics data

Over Representation Analysis was performed on the up-regulated proteins and genes from the bulk proteomics and bulk transcriptomics statistical results, respectively. For this purpose, clusterProfiler R package (v.4.14.6) was used (compareCluster function, minGS and maxGS was set at 15 and 500 respectively) along with ReactomePA (v1.50.0) and org.Hs.eg.db (v3.20.0) with a p-value and q-value cutoff set at 5%. All figures were created with tidyverse (v2.0) and more specifically ggplot2 (v3.5.2). For specific purposes such as heatmap and network enrichment plot, ComplexHeatmap (v2.22.0) and enrichplot (v1.29.2.003) R packages were used, respectively.

### Single-cell transcriptomics

#### Sample preparation

5 mm full-thickness skin biopsies were obtained from healthy donors and from Hailey-Hailey patients from both lesional and non-lesional areas. Immediately after collection, biopsies were rinsed with 50 µg/mL of gentamicin (Sigma-Aldrich, ref: G1397) diluted in Hanks’ Balanced Salt Solution (HBSS) (Gibco, ref: 14170-088). Biopsies were then incubated overnight at 4 °C in 1 U/mL of Dispase II/HBSS (Sigma-Aldrich, ref: D4693), after which the epidermis was peeled from the dermis. The two compartments were then processed separately.

#### Dermal single-cell dissociation

The dermis was cut into 1mm^2^ fragments and were digested for 3 hours at 37 °C with orbital shaking in enzyme cocktail using the Whole Skin Dissociation Kit, human (Miltenyi Biotec, ref: 130-101-540) according to the manufacturer’s instructions. Enzymatic activity was quenched by adding 10% FBS/DMEM (Sigma-Aldrich, ref: C8056 and Gibco, ref: 31966-021). The suspension was sequentially filtered through 70 µm and 40 µm cell strainers (Corning, ref: CLS431751 and CLS431750, respectively).

#### Epidermal single-cell dissociation

The epidermis was cut into 1mm^2^ fragments and incubated in 0.05% trypsin-EDTA (Gibco, ref: 25300-054) for 1 hour at 37 °C with manual agitation every 15 minutes. Digestion was quenched with 10% FBS/DMEM (Eurobio-Scientific, ref: CVFSVF06-01), and the cell suspension was sequentially filtered through 70 µm and 40 µm strainers.

#### Library preparation, sequencing, and alignment

Epidermal and dermal single-cell suspensions were counted and combined at a 1:1 ratio prior to library construction. Libraries were prepared at the Imagine Institute Genomics Platform (Paris, France) using the 10x Genomics Chromium Next GEM (v3.1) and GEM-X (v4) Single Cell 3′ Reagent Kits. Libraries were sequenced on Illumina NovaSeq 6000 using S2 flow cell with a 28-10-10-90 cycle configuration (paired-end: R1 = 28 bp, i7 = 10 bp, i5 = 10 bp, R2 = 90 bp). Sequencing reads were demultiplexed and aligned to the human reference transcriptome (GRCh38-2020-A from 10x Genomics).

#### Quality control and integration

Filtered UMI count matrices generated by Cell Ranger were analyzed using *Seurat* (v5.0.2) ^76^. Low-quality cells were excluded by filtering out barcodes with fewer than 500 detected genes and cells with >20% mitochondrial transcripts. After normalization, 3,000 highly variable genes were selected for downstream analyses. Putative heterotypic doublets were then identified and removed using *scDblFinder* (v1.24.0) ^77^, applying a stringent setting based on 3,000 top features. To integrate samples across conditions, we used *Seurat*’s anchor-based reciprocal PCA (RPCA) workflow. Integrated data were scaled, PCA was performed using the 3,000 variable genes, and UMAP was computed from the top 30 principal components. Clustering was performed by constructing a shared nearest-neighbor graph and applying the Louvain community detection algorithm. The optimal clustering resolution was selected by comparing clustering solutions across multiple resolutions using *clustree* (v0.5.1) ^78^; resolution 1 was retained for downstream analyses.

#### Annotation and downstream analysis

Cell type labels were assigned to the clusters based on a curated list of markers and cell types (**Supplementary Figure S3**). Clusters that featured markers of multiple different major cell types, high mitochondrial content and low number of features were considered contamination and subsequently removed from the integration. To validate cell type annotations, top genes expressed across all clusters were assessed using Seurat’s *FindAllMarkers* function.

UMAP projections and percentage cell proportion plots were generated using R package *dittoSeq* (v1.22.0). *Pheatmap* (v1.0.13) for the heatmap generated from aggregated and normalised (pseudobulk) cell counts and *CellChat* (v1.6.1) for net plot and activity bar plots.

#### Visium SD spatial transcriptomics

##### Sample processing, library preparation, and sequencing

Full-thickness skin biopsies were snap-frozen in isopentane/liquid nitrogen and embedded in OCT compound. Samples were sectioned at 12 µm and mounted onto Visium Spatial Gene Expression slides (10x Genomics). Subsequently, methanol fixation and H&E staining were performed according to the Visium Spatial Protocols (CG000240 and CG000160). Libraries were generated at the Imagine Institute Genomics Platform (Paris, France) using Visium Spatial Gene Expression 3’ reagent kit (10x Genomics). Sequencing was conducted on Illumina NovaSeq 6000 using an S2 flow cell with a 28-10-10-90 cycle configuration (paired-end: R1 = 28 bp, i7 = 10 bp, i5 = 10 bp, R2 = 90 bp).

##### Raw data processing and contamination removal

Visium data were loaded into the R environment with *Seurat* v5. Raw HDF5 matrices were additionally imported for contamination estimation. Spatial decontamination was performed using *SpotClean*, using 10 optimization iterations and a candidate radius of 20 pixels. Pre-and post-cleaning signals were evaluated using marker genes and spatial heatmaps. Decontaminated counts were converted back into a *Seurat* object with spatial image registration preserved. Contamination rates were retained in metadata.

##### Quality control and outlier removal

Spots were annotated with mitochondrial percentage, library complexity, and total/feature counts. Genes expressed in ≤3 spots were removed. Spot-level QC used median absolute deviation (MAD)-based detection of outliers for both total counts and total features in raw and log-transformed space. Spots failing log-MAD low thresholds for either counts or features were removed, along with spots exceeding 10% mitochondrial RNA.

##### Normalization, dimensionality reduction, and clustering

Filtered objects were normalized using *LogNormalize*. Ribosomal gene content (RP[SL]) was quantified per spot and regressed during scaling. PCA was performed using 100 components. Variance explained across components was calculated to guide downstream analyses. Graph-based clustering was performed across multiple resolutions (0-1.6) and cluster stability was evaluated using *clustree*; the resolution with maximal stability was selected for final clustering. UMAP embeddings were computed using the leading principal components.

##### scRNA-to-Visium label transfer and spatial cell type mapping

To annotate Visium spots with cellular identities, we used *Seurat* anchor-based transfer. After normalizing the Visium spatial assay and identifying variable genes, we intersected these with variable genes from the scRNA-seq reference and performed PCA on the Visium data (30 PCs). Transfer anchors were generated using *FindTransferAnchors* with log-normalization and 30 dimensions. Fine-grained and coarse-grained annotations were transferred separately. For each spot, the predicted cell type was defined as the class with maximal transferred probability; prediction confidence was defined as the maximum probability across labels. Shannon entropy of predicted probability vectors was computed per spot to quantify uncertainty. Cell type composition per spot was visualized using scatterpie representations overlaying spatial coordinates. Minor cell type probabilities were also projected as continuous features.

##### UCell-based IFN and IL-4/IL-13 pathway scoring in Visium

To quantify spatial interferon activation, we applied *UCell* scoring to Visium spots using an extended IFN/Reactome signature (IFN_REACTOME), which includes ISGs, JAK-STAT components, HLA class I/II genes, TRIM antiviral family members and nuclear pore translocation factors. *UCell* scores were computed for each spot and smoothed using *SmoothKNN* in PCA space to generate spatially coherent expression gradients. Smoothed were visualized using *SpatialFeaturePlot* with fixed cutoffs across samples. A similar workflow was applied to a curated IL-4/IL-13 inflammatory gene set (IL4_13_REACTOME), representing Th2 signaling and downstream response genes, yielding spatial IL-4/IL-13 activity maps.

### Xenium in situ transcriptomics

#### Sample processing and in situ profiling

Full-thickness skin biopsies were fixed in formalin, dehydrated through graded alcohol, and paraffin-embedded. The blocks were subjected to Xenium in situ transcriptomics preparation and sequencing at Advanced Genomics Core (University of Michigan, USA).

#### Data processing

Raw Xenium data were imported into Seurat and subjected to standard quality control. Cells were filtered based on transcript count (>100), gene number (>100), probe-based quality metrics (blank, control, and genomic-control fractions <0.20), and mitochondrial RNA content (<25%, when available). Filtered cells were normalized, and 2,000 highly variable genes were selected. Data was scaled, followed by principal component analysis (PCA), nearest-neighbor graph construction and clustering using the Louvain algorithm (resolution 0.2). UMAP embeddings were computed using the top principal components. Cell types were annotated by label transfer from the scRNA-seq dataset using Seurat’s reference-based mapping approach, leveraging shared gene expression profiles between modalities.

#### Spatial definition of epidermal compartments and immune infiltration

To quantify immune localization relative to the dermal–epidermal junction (DEJ), a spline-based model was fitted to the spatial coordinates of basal keratinocytes in each sample. This curve was used to approximate the DEJ and classify cells as epidermal or dermal based on their relative position.

#### Cell-cell contact analysis from Xenium coordinates

Cell-cell proximity was quantified by computing Euclidean pairwise distances between all cells and defining a contact as <50 µm. Contact adjacency matrices were computed separately for control and lesional samples. Matrices were aligned to a shared set of cell types, and differential contact frequency was computed as the difference in adjacency entries between lesion and control. To assess statistical significance, a permutation test was conducted in which total contacts for each cell type pair were redistributed between conditions using a binomial model. For each pair, 1,000 permutations were generated, and empirical two-sided P values were computed as the fraction of permuted differences exceeding the observed absolute difference. P values were corrected using FDR. Differential contacts were visualized using *ComplexHeatmap* with classes grouped as epithelial, immune, stromal, endothelial, and vascular.

### Drug repurposing analysis

#### Interaction-based target-drug map

The nCounter Human Immunology V2 Panel gene list (Bruker Spatial Biology) was used as the input gene set. Each input gene was used to query the Open Targets Platform GraphQL API ^40^ to retrieve molecular interaction partners from the Reactome ^79^ and STRING ^80^ databases. To filter out STRING low-confidence interactions, the database score and overall interaction score thresholds were set to ≥0.4 and ≥0.7, respectively. Partners from the two databases were then merged and deduplicated. Input genes and their interaction partners were subsequently queried against the Open Targets Platform GraphQL API for linked approved drugs to generate an integrated target-drug map.

#### Immune-related module scoring

Immune-related modules for type I interferon, eosinophilic, macrophagic, neutrophilic, Th1, Th2, and Th17, as established by Seremet, T. *et al.*^38^, and IFN-γ (type II interferon), as established by Ayers, M. *et al.* ^39^, were applied to the bulk transcriptomic data. For each sample, the score for a given module was calculated as the mean variance-stabilized expression of all genes assigned to that module. A module was considered activated when its score exceeded the threshold defined as the mean score of non-lesional samples + 2 s.d.

#### Candidate drugs selection

Differential gene expression analysis between lesional and non-lesional samples was performed using DESeq2 ^81^. Genes within the immune-related modules that met an adjusted *P* value < 0.05 were queried against the integrated target-drug map. Upregulated genes were matched to inhibitors or antagonists, whereas downregulated genes were matched to agonists or activators. Candidate drugs were ranked according to the number of matched targets. Final drug selection criteria were: (1) suppression of IFN signaling, (2) broad coverage of dysregulated immune modules, and (3) suitability for topical application.

#### Blood IFN score

Total RNA was extracted from whole blood of HH patients using the PAXgene Blood RNA Kit (PreAnalytiX, ref: 762164), and analysis was conducted at the Centre de Biologie et de Pathologie Sud (Hospices Civils de Lyon, France). The interferon score was determined by measuring the expression of six interferon-stimulated genes (*IFIT1*, *ISG15*, *RSAD2*, *IFI27*, *SIGLEC1*, and *IFI44L*) using the nCounter system (NanoString). Data was normalized to internal housekeeping genes and compared with a healthy control reference population. A threshold score of > 2.3 was used to define a positive type I IFN signature.

#### Outcomes and clinical measures

Patient-reported outcomes (PROs) were collected at baseline (week 0) and at subsequent, clinically determined follow-up visits. Dermatology Life Quality Index (DLQI): a 10-item questionnaire assessing quality of life (QoL) with a total score range of 0-30 (where 0-1 = no effect; 2-5 = small; 6-10 = moderate; 11-20 = very large; 21-30 = extremely large effect on QoL) ^82^. Skindex-16: a 16-item, dermatology-specific QoL instrument; total score 0-96 (higher scores indicate worse quality of life) ^83^. Numeric Rating Scales (NRS): 11-point scales (0-10) for average itch and average pain (where 0 = no symptoms, 1-3 = mild, 4-6 = moderate, 7-8 = severe, and 9-10 = very severe).

To standardize outcomes across variable visits, one acute period and follow-up assessment periods were specified relative to baseline (Early: days 1-14; Month 1: days 15-44; Month 2: days 45-74; Month 3: days 75-104; Month 4: days 105-134; Month 5: days 135-164; Month 6: days 165-194; Month 7: days 195-224; Month 8: days 225-254; Month 9: days 255-284). When multiple reports were available for a patient within the same assessment period, the report closest to the predefined target day was selected (day 7 for Early, day 30 for Month 1, day 60 for Month 2, day 90 for Month 3, and so on). Patients without an available report within a given period were excluded from descriptive summaries for that period. Clinical outcomes data are presented in the result as medians [minimum-maximum].

### Histological analysis and Immunostaining

Full-thickness skin biopsies were fixed in formalin, dehydrated through graded alcohol, and paraffin-embedded. Paraffin blocks were sectioned at 5 µm and used for hematoxylin and eosin (H&E) or immunofluorescence staining. For H&E staining, slides were automatically deparaffinized, rehydrated, stained using Leica ST5020-CV5030 integrated stainer workstation (Leica Biosystems, France), and scanned using NanoZoomer S60 (Hamamatsu Photonics, Japan). For immunofluorescence, sections were deparaffinized, rehydrated, and subjected to heat-induced epitope retrieval using citrate buffer (pH 6.0) or Tris-EDTA buffer (pH 9.0; for STAT1 and pSTAT1 staining). The following primary antibodies were used: pSTAT1 (Cell Signaling Technology, ref: 9167), pSTAT3 (Cell Signaling Technology, ref: 9145), IRF9 (Proteintech, ref: 14167-1-AP), IDO1 (Invitrogen, ref: 14-9750-82), BDCA2 (R&D Systems, ref: AF1376), CD3 (Abcam, ref: ab11089), CD8 (Abcam, ref: ab101500), CD14 (Abcam, ref: ab183322), CD20 (Abcam, ref: ab64088), CD68 (Abcam, ref: ab213363), CD207 (Atlas Antibodies, ref: HPA011216), NCAM1 (CD56) (Abcam, ref: ab75813), MPO (R&D Systems, ref: MAB3174), Loricrin (BioLegend, ref: PRB-145P), Filaggrin (BioLegend, ref: PRB-417P), KRT1 (Sigma-Aldrich, ref: HPA017917), KRT14 (Leica Biosystems, ref: NCL-L-LL002) and KI67 (Invitrogen, ref: 740008T). Secondary antibodies were: Alexa Fluor™ Plus 488-conjugated goat anti-mouse IgG (Invitrogen, ref: A32723), Alexa Fluor™ Plus 555-conjugated goat anti-rabbit IgG (Invitrogen, ref: A32732), Alexa Fluor™ Plus 647-conjugated goat anti-rat IgG (Invitrogen, ref: A48265), and Alexa Fluor™ Plus 555-conjugated donkey anti-goat IgG (Invitrogen, ref: A32816). Slides were mounted using Fluoromount-G™ with DAPI (Invitrogen, ref: 00-4959-52) to counterstain nuclei. Images were acquired on Leica TCS SP8 SMD confocal microscope (Leica Microsystems, France) and processed using ImageJ (NIH, USA). Images shown are representative of at least three biological replicates. Graphs and statistical analyses were performed using GraphPad Prism (v9.1.0).

## Data availability

Supplementary Tables and multiomics data will be available upon publication. Multiomics data will be available via Zenodo. The mass spectrometry proteomics data have been deposited to the ProteomeXchange Consortium via the PRIDE^84^ partner repository with the dataset identifier PXD078090 and will be publicly available following publication.

## Acknowledgements

The authors are grateful to the HHD patients who participated in this study, to the Association pour les maladies de Hailey-Hailey et Darier (AFRHAIDA), to the Association Ichtyose France (AIF) and to the French Society of Dermatology (Société Française de Dermatologie, SFD). We thank Lola Taveau for technical help, Mickaël Ménager and Marine Luka from the LabTech Single-Cell@Imagine of the Institute Imagine. A.H. received support from ANR (ENIGMncA, ANR-21-CE14-0040-02 grant). J.E.G. was supported by the NIH-P30-AR075043 grant.

## Contributions

S.C. performed proteomics analyses, critically analyzed and interpretated omics datasets, and wrote the manuscript. H.D.M.P. performed scRNA-seq analyses, supervised immunostaining experiments, analyzed patient metadata, and wrote the manuscript. M.D. performed immunostaining experiments. J.M. analyzed the scRNA-seq dataset. K.R. integrated bulk transcriptomic and proteomic data. L.C.T., P.H., and J.E.G. generated spatial transcriptomic data.

J.E.G. critically reviewed the manuscript and provided resources. M.E.P.L. analyzed scRNA-seq and spatial transcriptomic datasets and supervised the study. I.G.C. supervised the study, provided resources, and critically reviewed the manuscript. A.H. conceived and supervised the study, collected patient samples, treated patients, provided resources, and wrote the manuscript. All authors discussed and approved the final version before submission.

## Ethics declaration

The authors declare no competing interests.

## Notes

### Competing Interest Statement

The authors have declared no competing interest.

