## Supplementary Figures for "Multiomic Analysis Reveals an IFN-driven Cellular Landscape Effectively Targeted by Ruxolitinib in Hailey-Hailey Disease"

### Figure S1

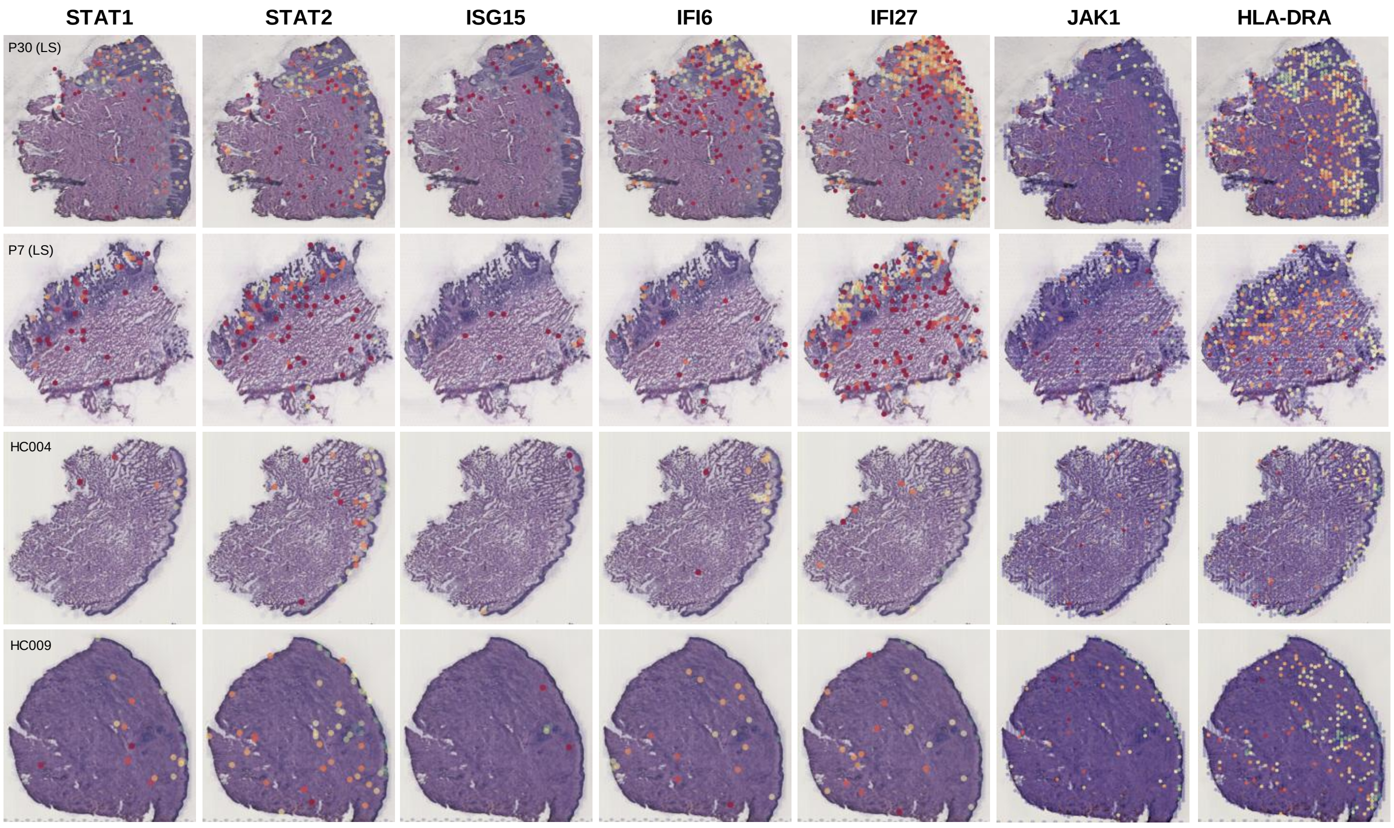


**Figure S1. Upregulation of IFN signaling in lesional skin in Hailey-Hailey disease.** Spatial plots showing the expression level of STAT1, STAT2, ISG15, IFI6, IFI27, JAK1, HLA-DRA in spatial-seq samples (VisiumSD assays).

#### Figure S2

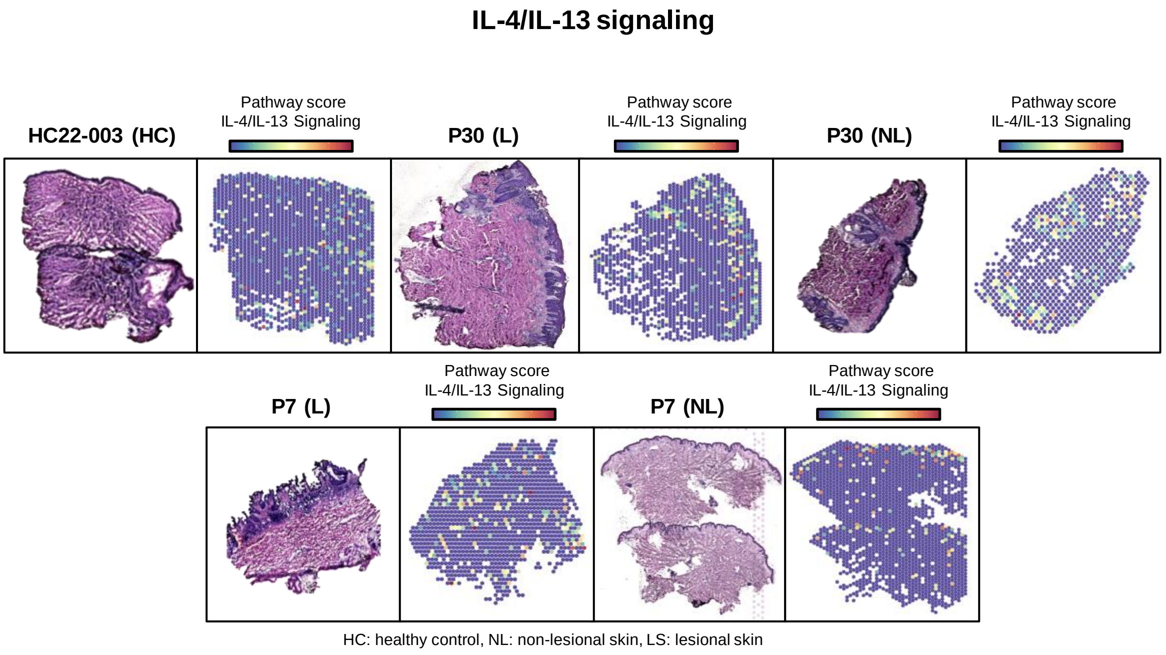


**Figure S2. Modest upregulation of IL-4/IL-13 signaling in lesional skin in Hailey-Hailey disease.** Spatial transcriptomics maps (VisiumSD assays) showing IL-4/IL-13 signaling pathway activity scores in representative healthy control (HC), patient non-lesional and lesional skin sections.

**Figure S3**

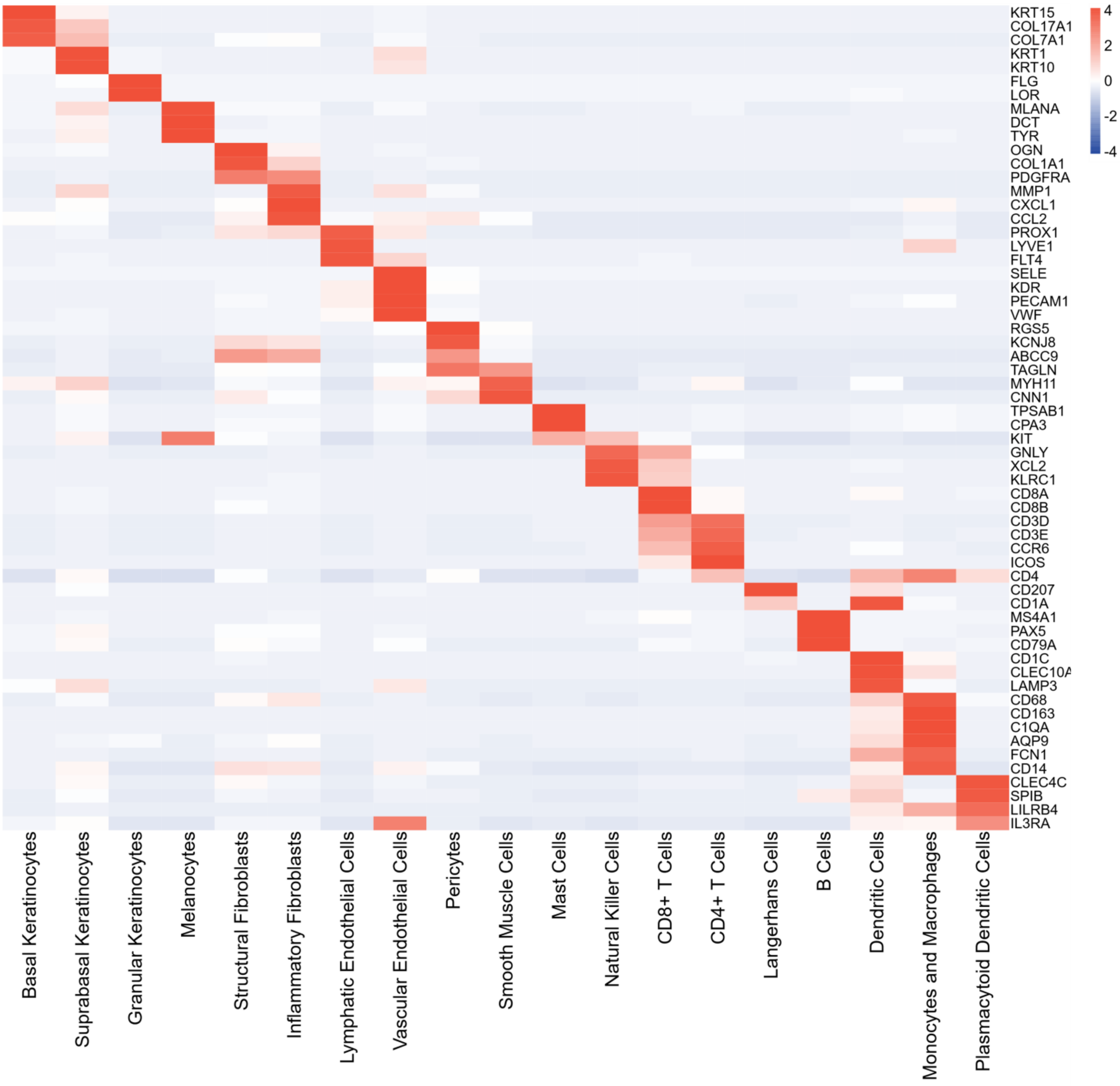


**Figure S3. Heatmap showing scRNA-seq expression profiles of annotated cell types.** The heatmap shows the expression of cell-type-specific markers used to identify cell populations in scRNA-seq data from 11 skin samples (4 HC, 3 NL, and 4 L). Columns represent identified cell-type clusters, and rows represent curated marker genes. The color scale represents the scaled expression of each gene.

#### Figure S4

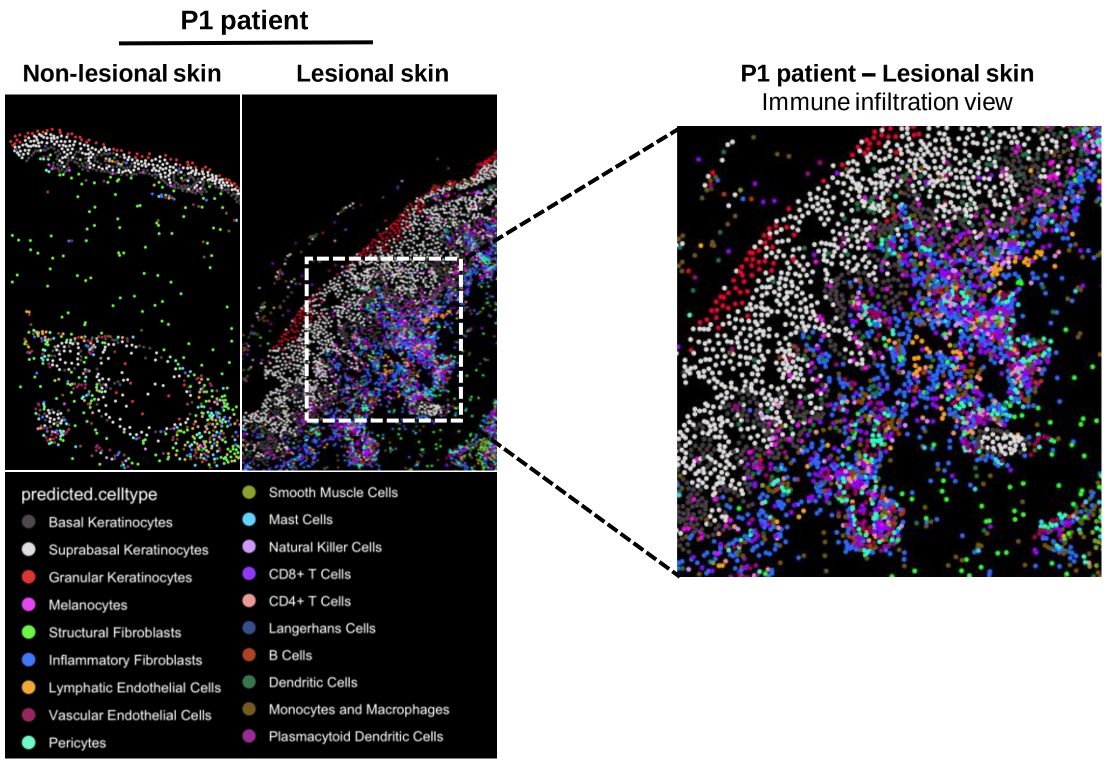


**Figure S4. Single-cell resolution mapping of immune infiltration in lesional skin.** Zoomed view on the region labeled in 3A by the white square, showing immune infiltration beneath and within the epidermal layers in the lesional sample.

**Figure S5**

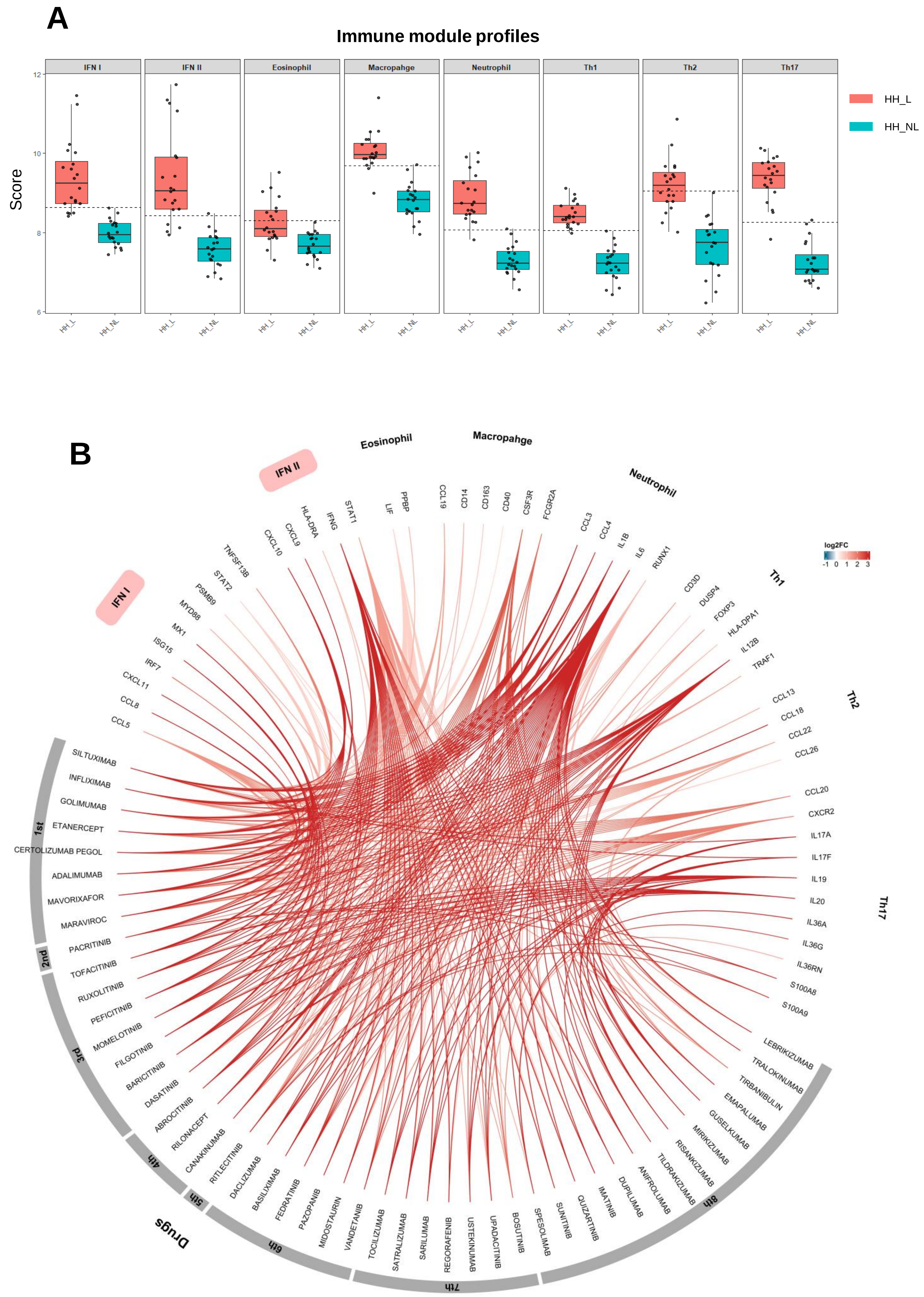


**Figure S5. Immune module activation profiles and molecular interaction-based drug repurposing analysis in Hailey-Hailey disease. (A)** Immune module scores of Hailey-Hailey lesional skin (HH_L: pink) and non-lesional skin (HH_NL: blue). Module scores were calculated from predefined immune gene sets. The black dashed line indicates the activation threshold, defined for each module as the mean + 2sd of the HH_NL reference distribution. Boxes represent the median and IQR, whiskers extend to 1.5 × IQR, and points indicate individual samples. **(B)** Circos plot showing the relationships between differentially expressed genes (DEGs) in Hailey-Hailey disease and candidate therapeutic agents. Upper segments represent immune modules (interferon, eosinophil, macrophage, neutrophil, Th1, Th2 and Th17) containing DEGs identified in lesional skin compared with non-lesional skin. The lower segment lists candidate repurposed drugs, ordered by the total number of gene targets (left to right, highest to lowest). Ribbon connections indicate target-drug interactions, with colors represent gene log2 fold change (red, upregulated; blue, downregulated). Only approved drugs targeting ≥5 DEGs are shown.

**Figure S6**

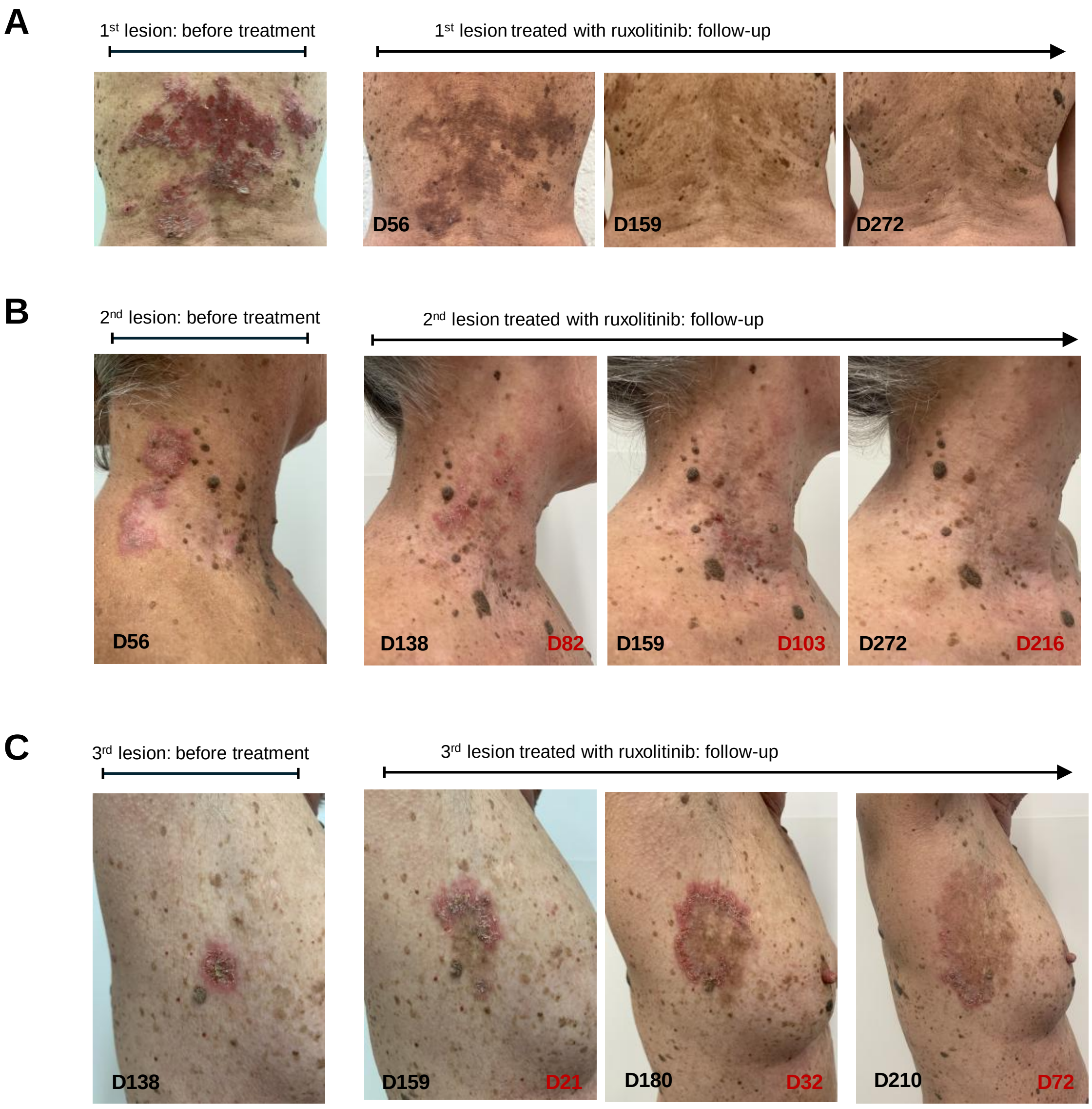


**Figure S6. Occurrence of new skin lesions and clinical improvement following ruxolitinib treatment in patient HH1. (A)** Representative clinical photographs of the first lesion treated with ruxolitinib. **(B)** A second lesion occurred 2 months after initiation of treatment of the first lesion, in an untreated area. **(C)** Representative of another lesion occurred at the periphery of the treated area. These newly occurring lesions responded well to ruxolitinib treatment. Day labels in black indicate time from initiation of ruxolitinib treatment of the first lesion. Day labels in red indicate time from initiation of ruxolitinib treatment of the corresponding lesion.

#### Figure S7

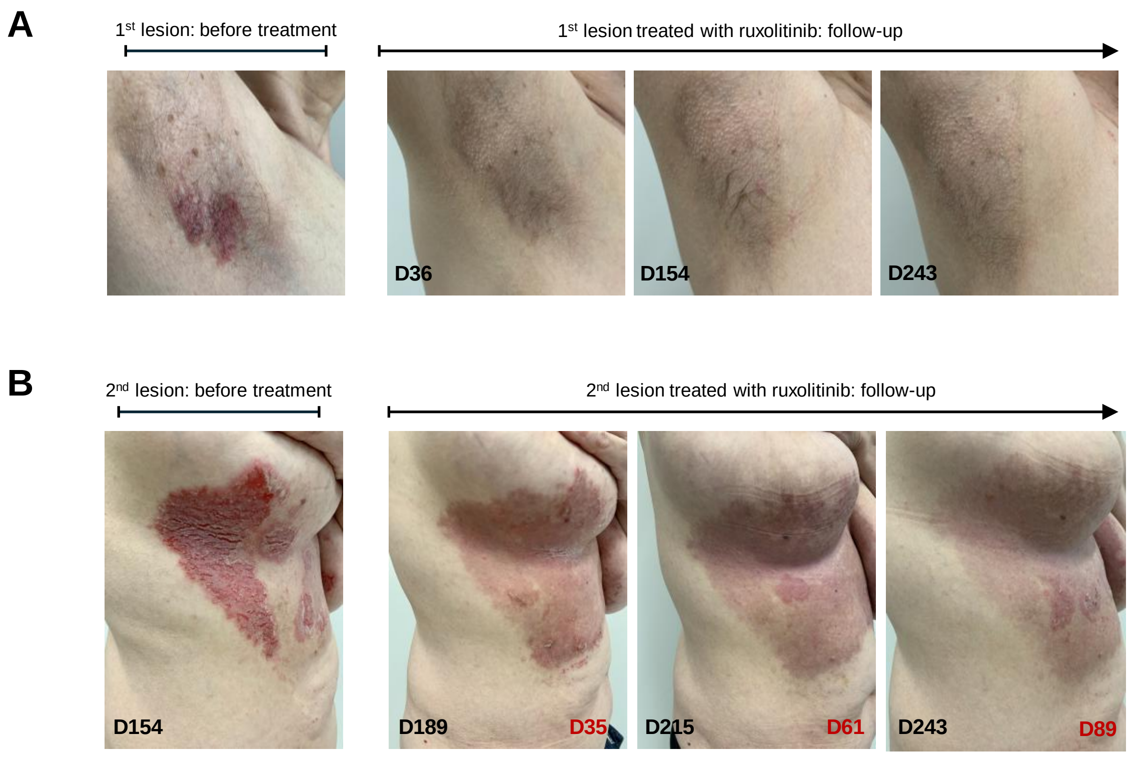


**Figure S7. Occurrence of new skin lesion and clinical improvement following ruxolitinib treatment in patient HH5. (A)** Representative clinical photographs of the first lesion treated with ruxolitinib. **(B)** A second lesion occurred 5 months after initiation of treatment of the first lesion, in an untreated area. The newly occurring lesion responded well to ruxolitinib treatment. Day labels in black indicate time from initiation of ruxolitinib treatment of the first lesion. Day labels in red indicate time from initiation of ruxolitinib treatment of the corresponding lesion.

#### Figure S8

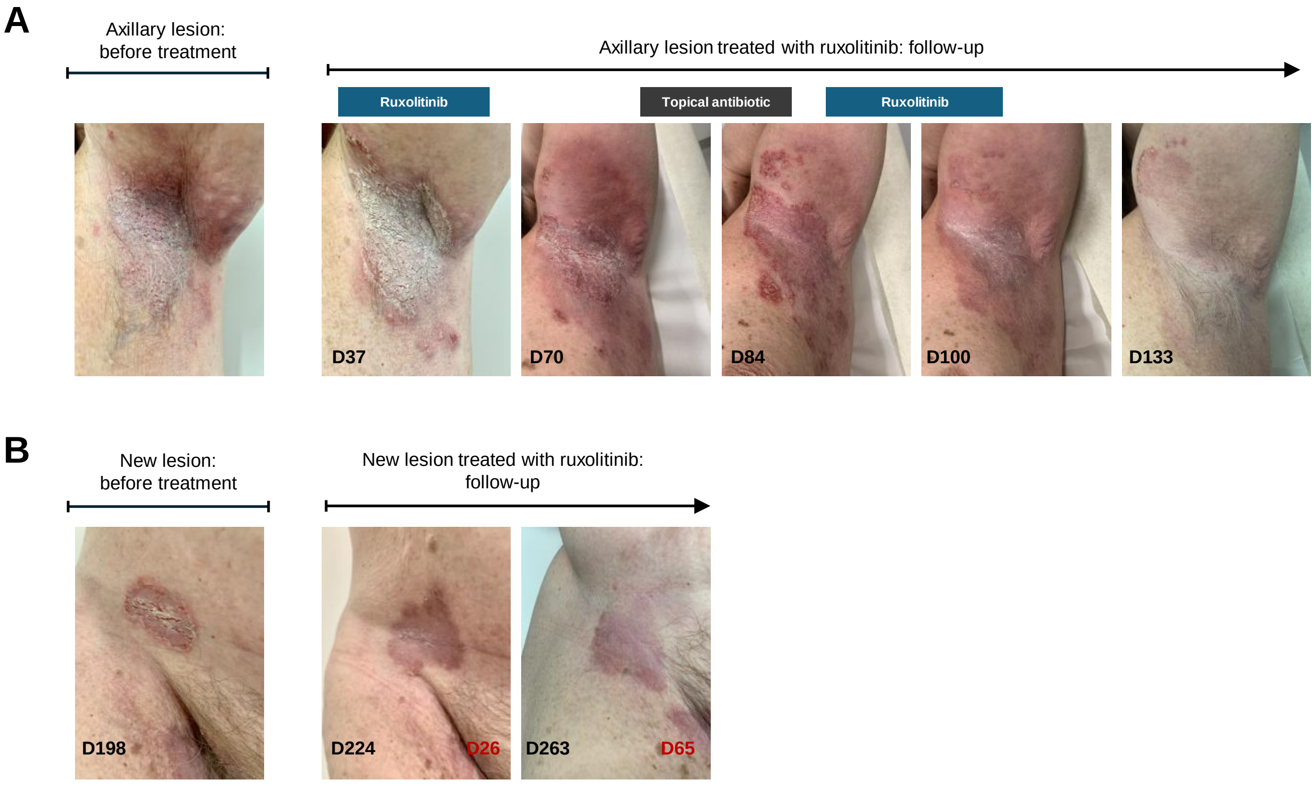


**Figure S8. Occurrence of a new skin lesion and clinical improvement after antibiotic treatment followed by ruxolitinib in patient HH2. (A)** Representative clinical photographs of the axillary lesion. The clinical course suggests that antibiotic treatment before initiation of ruxolitinib may be required. **(B)** A new lesion occurred 7 months after initiation of treatment of the first lesion, in an untreated area. The newly occurring lesion responded well to ruxolitinib treatment. Day labels in black indicate time from initiation of ruxolitinib treatment of the first lesion. Day labels in red indicate time from initiation of ruxolitinib treatment of the corresponding lesion.

Figure S9

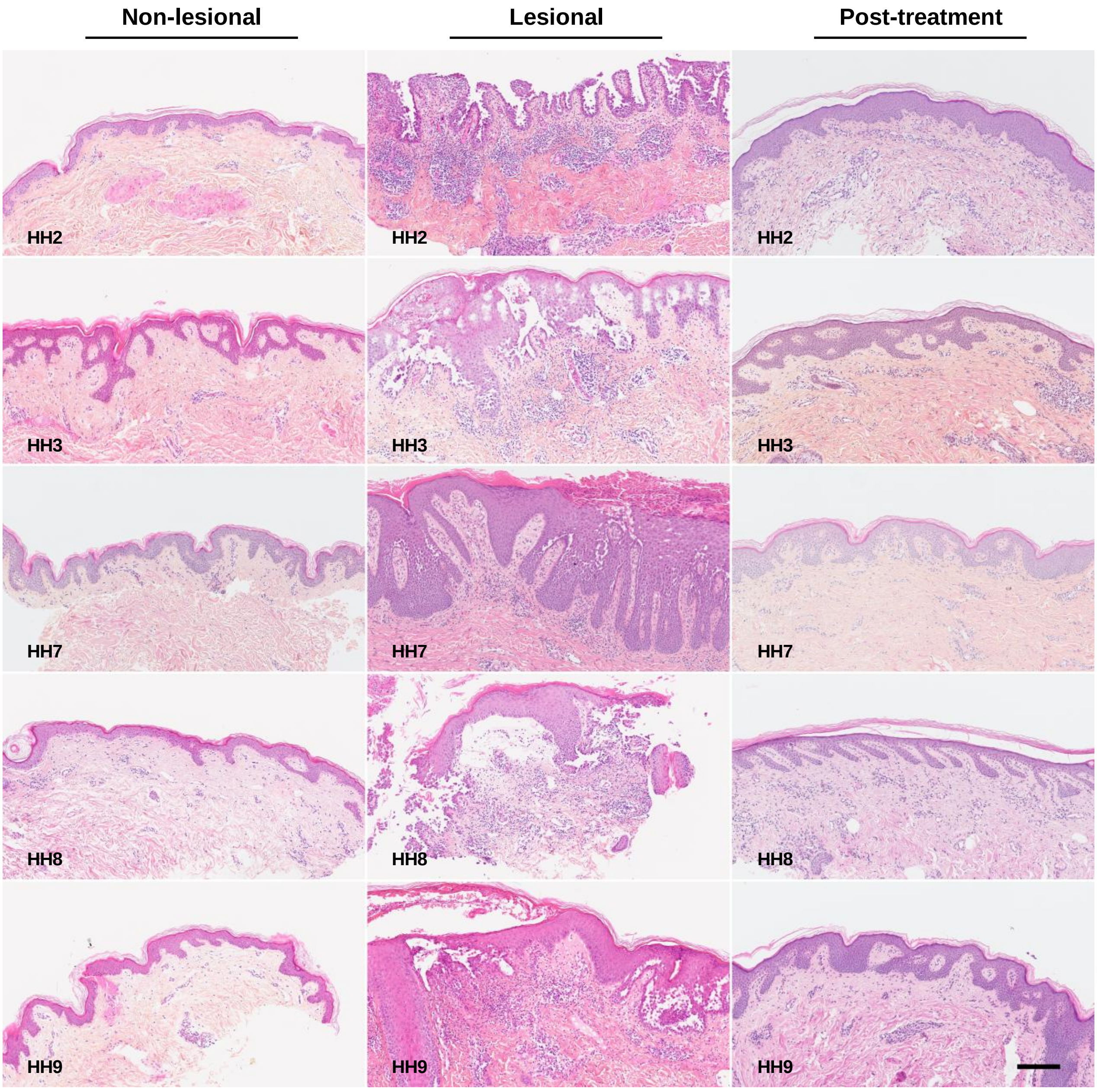


**Figure S9. Clinical and histopathological improvement of Hailey-Hailey (HH) lesional skin following ruxolitinib treatment.** Haematoxylin and eosin stained sections of non-lesional, lesional and post-treatment (PT) skin from HH patients. PTL skin for HH2, HH3, HH7, HH8 and HH9 were biopsied at days 69, 35, 69, 36 and 32 after ruxolitinib treatment initiation, respectively. Scale bar = 200 µm.

### Figure S10

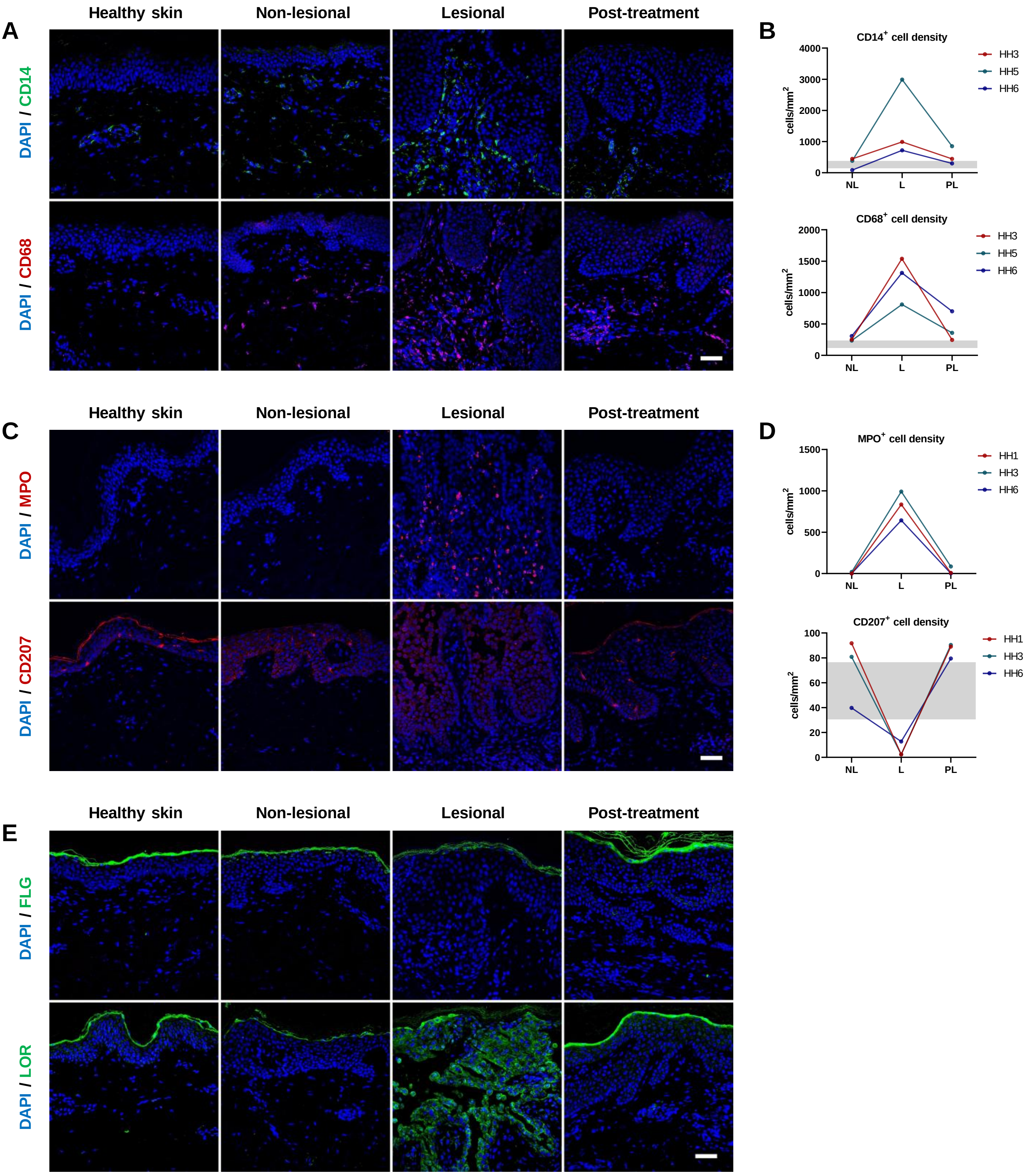


**Figure S10. Ruxolitinib broadly reduces the cutaneous immune cell infiltrate and restores epidermal homeostasis in Hailey-Hailey disease. (A,C)** Representative immunofluorescence images of innate immune cell populations, including **(A)** monocytes (CD14, green) and macrophages (CD68, red), and **(C)** neutrophils (MPO, red) and Langerhans cells (CD207, red). **(B,D)** Quantification of cell densities (cells per $mm^2$) for the populations described in A-D across non-lesional (NL), lesional (L), and post-treatment (PT) skins. Gray shaded areas represent the range measured in healthy controls (n=3). **(E)** Representative immunofluorescence images of the barrier proteins filaggrin (FLG, green) and loricrin (LOR, green). Nuclei were counterstained with DAPI (blue). Scale bars = 50 µm. Data points represent individual patients (n=3).

**Figure S11**

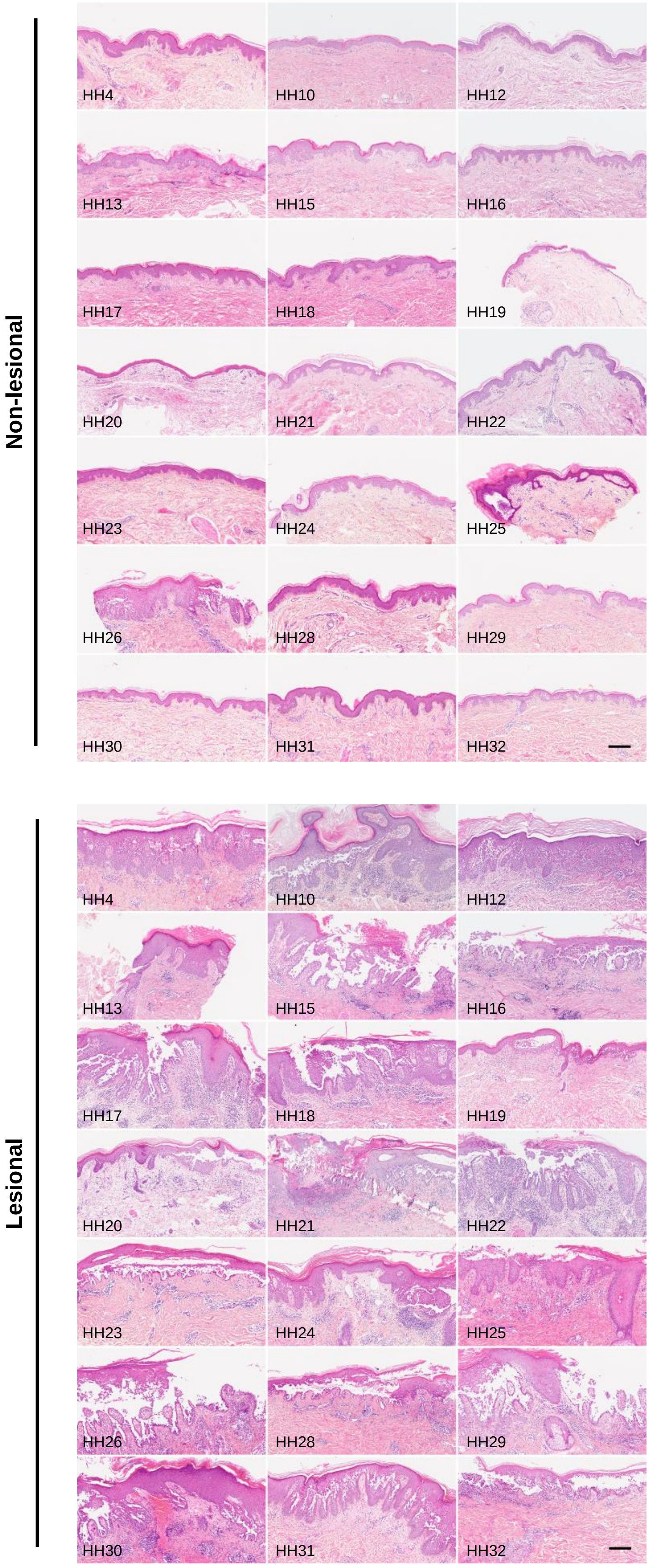


**Figure S11. Histopathology of Hailey-Hailey (HH) skin.** Non-lesional and lesional skin sections from HH patients stained with haematoxylin and eosin. Scale bar = 200 µm.
